# Insights from aquaporin structures into drug-resistant sleeping sickness

**DOI:** 10.1101/2025.05.15.654314

**Authors:** Modestas Matusevicius, Robin A. Corey, Marcos Gragera, Keitaro Yamashita, Teresa Sprenger, Marzuq A. Ungogo, James N. Blaza, Pablo Castro-Hartmann, Dimitri Y. Chirgadze, Sundeep Chaitanya Vedithi, Pavel Afanasyev, Roberto Melero, Rangana Warshamanage, Anastasiia Gusach, Jose Maria Carazo, Mark Carrington, Tom L. Blundell, Garib Murshudov, Phillip Stansfeld, Mark Sansom, Harry P. de Koning, Christopher G. Tate, Simone N. Weyand

## Abstract

Trypanosoma brucei is the causal agent of African trypanosomiasis in humans and animals, the latter resulting in significant negative economic impacts in afflicted areas of the world. Resistance has arisen to the trypanocidal drugs pentamidine and melarsoprol through mutations in the aquaglyceroporin TbAQP2 that prevent their uptake. Here we use cryogenic electron microscopy to determine the structure of TbAQP2 from Trypanosoma brucei, bound to either the substrate glycerol or to the sleeping sickness drugs, pentamidine or melarsoprol. The drugs bind within the AQP2 channel at a site completely overlapping that of glycerol. Mutations leading to a drug-resistant phenotype were found in the channel lining. Molecular dynamics simulations showed the channel can be traversed by pentamidine, with a low energy binding site at the centre of the channel, flanked by regions of high energy association at the extracellular and intracellular ends. Drug-resistant TbAQP2 mutants are still predicted to bind pentamidine, but the much weaker binding in the centre of the channel observed in the MD simulations would be insufficient to compensate for the high energy processes of ingress and egress, hence impairing transport at pharmacologically relevant concentrations. The structures of drug-bound TbAQP2 represent a novel paradigm for drug-transporter interactions that could provide new mechanisms for targeting drugs into pathogens and human cells.

## Introduction

Human African trypanosomiasis (HAT) remains a neglected tropical disease^1^. It is transmitted by tsetse flies and, once symptoms are apparent, treatment is necessary to avoid fatality. The infection starts with a haemolymphatic stage that progresses to a late stage infection of the central nervous system, at which point severe neurological symptoms occur with rapid progression to coma and death^2^. Early-stage infections are commonly treated with pentamidine and the late stage with melarsoprol, although for most cases this is now being replaced with fexinidazole^3^. Animal African trypanosomiasis (AAT) causes billions of dollars in damage to agriculture throughout the tropics^4,5^. For both HAT and AAT, chemotherapy remains the only control option and most of the medication is old, inadequate and threatened by resistance^3,4,6,7^.

For most trypanocidal drugs, resistance is associated with mutations in the membrane transporters through which the drugs enter the trypanosome^8^. The uptake of pentamidine and melarsoprol was initially attributed to a High Affinity Pentamidine Transporter (HAPT)^9,10^, which was subsequently identified as the aquaglyceroporin TbAQP2^11,12^. Deletions and chimeric rearrangements between TbAQP2 and the adjacent gene TbAQP3 were found to be responsible for melarsoprol-pentamidine cross-resistance (MPXR; Fig. S1)^6,12–15^. To understand the differences between TbAQP2 (water/glycerol/drug transporter) and TbAQP3 (classical aquaglyceroporin transporting water/glycerol^16^ but not pentamidine or melarsoprol^11,12,17^), mutants were made exchanging amino acids between the proteins^17^. Mutation of TbAQP2 to add any of the TbAQP3 selectivity filter residues Tyr250, Trp102 or Arg256 resulted in inhibition of melarsoprol and pentamidine uptake, whereas TbAQP3 containing all three of their TbAQP2 counterparts (Leu250, Ile102, Leu256) permitted the uptake of pentamidine. However, the molecular details of the interactions between TbAQP2 and drugs as well as the resistance mechanism remained unclear, with alternative models proposing that TbAQP2 acts as a channel^8,17,18^ or receptor^19^ for the drugs. To resolve this quandary and to resolve the molecular interactions underlying drug uptake by aquaporins, we determined the structures of TbAQP2 bound to either melarsoprol, pentamidine or the native substrate, glycerol.

### Cryo-EM structures of TbAQP2 and drug binding

TbAQP2 was expressed using the baculovirus expression system in insect cells and purified in the presence of either a drug (melarsoprol or pentamidine) or the substrate glycerol (see Methods, Fig. S2a,b). Structures of TbAQP2 bound to either glycerol, melarsoprol, or pentamidine were determined by electron cryogenic microscopy (cryo-EM) to overall resolutions of 3.2 Å, 3.2 Å and 3.7 Å, respectively (Figs. S2c, S3 and S4a-c). The resolution was sufficient to see the majority of the side chains and the bound drug or glycerol (Figs S5-S8). The cryo-EM structures of TbAQP2 (excluding the drugs/substrates) were virtually identical (rmsd 0.2 Å) and represented the aquaporin tetramer (Fig. 1a,b) that is typical for this family of channels^20^. The tetramer contained four channels, each consisting of a single TbAQP2 polypeptide folded into a bundle of eight α-helices (Fig 1b), two of which (helices H3 and H7) penetrated the plasma membrane only partially, as is typical for aquaporins^20^, the others crossing the membrane fully.

**Fig. 1.**
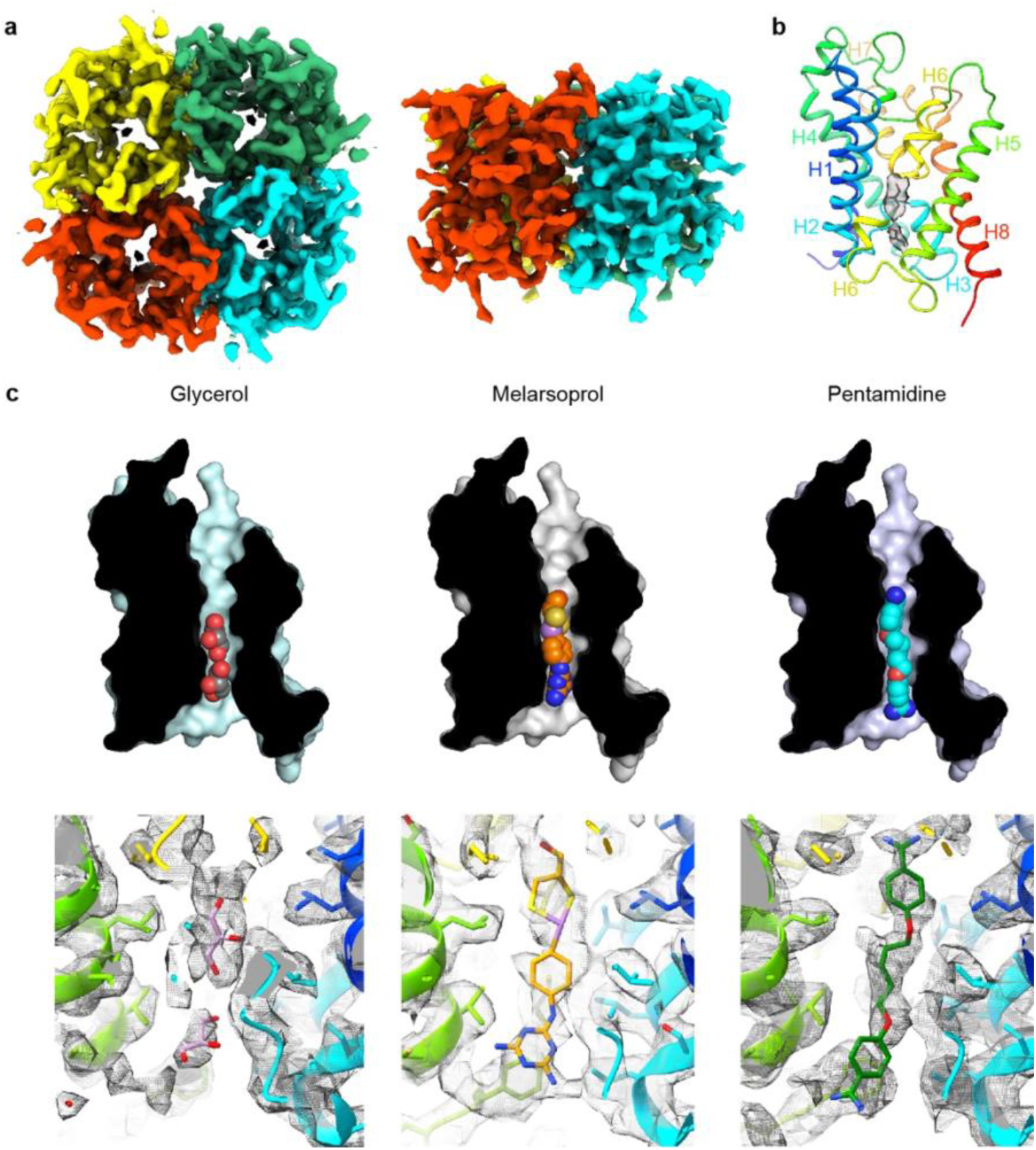
Cryo-EM structures of TbAQP2 bound to either glycerol, melarsoprol or pentamidine. **a**, Overall structure of the TbAQP2 tetramer viewed either from the extracellular surface or within the membrane plane. **b**, Structure of protomer A of TbAQP2 viewed as a cartoon (rainbow colouration) with glycerol (sticks) bound. The cryo-EM density for glycerol is shown as a grey surface. **c**, Cut-away views of the channel in each of the TbAQP2 structures showing bound substrates and drugs (spheres) with atoms coloured according to type: red, oxygen; yellow, sulphur; blue, nitrogen; purple, arsenic; carbon, grey (glycerol), orange (melarsoprol) or cyan (pentamidine). **d**, Cryo-EM densities (grey surface) for glycerol, melarsoprol and pentamidine in their respective structures. See Figures S7 and S8 for different views of the substrates and comparisons between densities.

Melarsoprol and pentamidine are bound in a channel formed by the TbAQP2 protomer, completely overlapping the binding site for the substrate glycerol (Fig. 1c,d). The densities of the substrates were distinct from one another and are consistent with the structures of the substrates (Figs. S7 and S8). The position of drug binding is consistent with a hotspot analysis performed on the structures (see Methods and Fig. S2d,e). This analysis defines likely regions where molecules may bind based on *in silico* docking scores of small molecule fragments docking to a model of TbAQP2^21^ (see Methods). Binding of the two glycerol molecules is mediated by a total of six hydrogen bonds, four to backbone carbonyl groups (residues Gly126, Gly127, His128, Leu129) and two to the side chain of His128 (Fig. 2a). These are on a polar strip of residues running down the channel, that also includes Val133, which makes van der Waals contacts with the glycerol molecule closest to the cytoplasm. The remainder of the contacts are van der Waals interactions (Val222, Phe226, Ile241, Val245) on the opposite side of the channel. It is striking that all eleven amino acid residues within 3.9 Å of the two glycerol molecules also make contacts to both pentamidine and melarsoprol (Fig. 2a). Whereas the TbAQP2-glycerol contacts are equally polar and non-polar, van der Waals interactions dominate contacts between the drugs and TbAQP2. The exceptions are a hydrogen bond between the backbone carbonyl of Ala259 and both drugs, and a backbone carbonyl hydrogen bond between Gly126 and melarsoprol that is also seen between glycerol and TbAQP2 (Fig. 2a). It is also remarkable that an overlay between the three different structures shows a very close alignment between glycerol, pentamidine and melarsoprol (Fig. 2b) which is consistent with the channel structure being virtually identical regardless of whether a drug or glycerol is bound. The size of the channel with pentamidine bound is very similar to the glycerol-bound *Plasmodium falciparum* AQP structure (Fig. S2f). The close fit of the drugs within the channel is consistent with the extremely limited modification of the drugs that is possible whilst retaining the ability to pass through the pore, including modifications that reduce flexibility^17^. Diamidines with fixed-angle structures such as diminazene and furamidine are not substrates for TbAQP2 but are exclusively accumulated through aminopurine transporter TbAT1^22,23^. A number of amino acid residues interact only with pentamidine and/or melarsoprol and not with glycerol (Leu84, Ile110, Val114, Leu122, Leu218, Ala259, Asn261, Met260, Leu264), which is a result of the drugs being longer and extending further along the channel towards the extracellular surface when compared with glycerol (Figs. 1d and 2b). Towards the intracellular surface, only Leu122 makes contacts to the drugs and not to glycerol. In all, 19 residues in TbAQP2 interact with pentamidine, compared with 18 residues interacting with melarsoprol and 11 to the two molecules of glycerol.

**Fig. 2.**
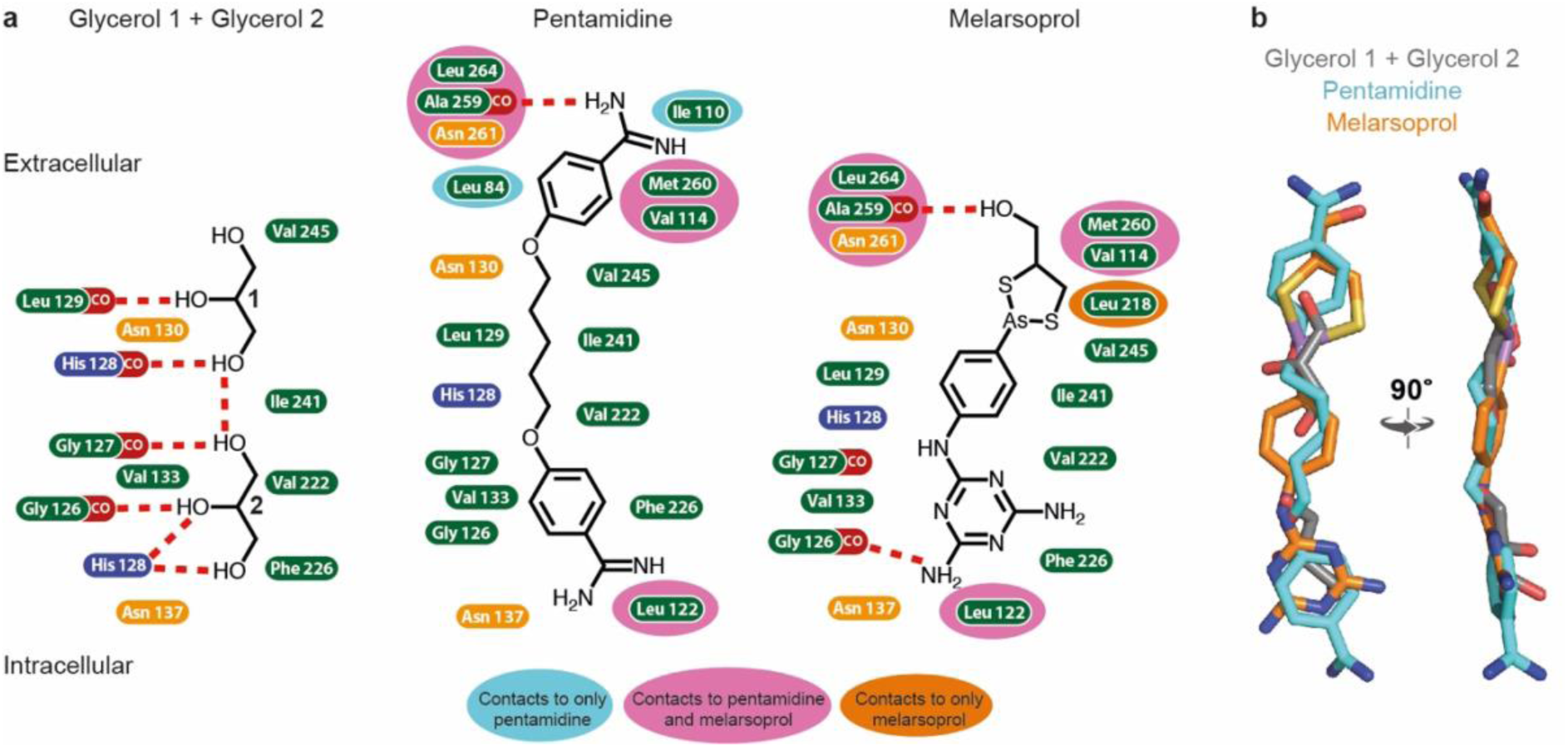
Interactions between TbAQP2 and bound substrates and drugs. **a**, Amino acid residues containing atoms ≤ 3.9 Å from substrate or drug model in the cryo-EM structures are depicted and coloured according to the chemistry of the side chain: hydrophobic, green, positively charged, blue; polar, orange. Interactions with a backbone carbonyl group is shown as CO (red) and potential hydrogen bonds are shown as red dashed lines. Residues highlighted in an additional colour (grey, pale blue, pink or dark orange) make contacts in two or less of the structures (as indicated on the figure), whilst those without additional highlighting make contacts in all three structures. **b**, Protomer A from each structure was aligned and the positions of the two glycerol molecules (grey), melarsoprol (orange) and pentamidine (cyan) are depicted.

The structure of AQP2 from *Plasmodium falciparum* (PfAQP)^24^ with glycerol bound is the closest structural homologue to TbAQP2 available (rmsd 0.8 Å, PDB ID 3CO2, 33% amino acid sequence identity). The overall arrangement of α-helices in TbAQP2 is identical to that in PfAQP (Fig. 3a-d) and other aquaglyceroporins whose structures have been determined (human AQP7^25^, human AQP10^26^ and *Escherichia coli* GlpF^27^). Despite the low identity in amino acid sequence, there are striking similarities between the structures of TbAQP2 and PfAQP in the conserved NPA/NPA motif and the positions of bound glycerol molecules in the channel. The NPA/NPA motif is found at the ends of the half-helices H3 and H7, with TbAQP2 and PfAQP similarly differing from the canonical sequences (NPS/NSA in TbAQP2 and NPS/NLA in PfAQP; Fig 3a,b). Superposition of TbAQP2 and PfAQP shows the position and rotamer of Asn-Pro-Ser in the NPS motif and both Asn and Ala in the NSA/NLA motifs are identical (Fig. 3c). Some of the network of hydrogen bonds that maintain the structure of this region are also conserved, such as the hydrogen bonds in TbAQP2 between Asn130 and the backbone carbonyl of 260 and the backbone amine of Ala132, the hydrogen bond between Ser263 and Gln155 and the hydrogen bond between Asn261 and the backbone amide of Ser263. However, both Asn residues in the NPS/NLA motif of PfAQP form hydrogen bonds with one of the glycerol molecules in the channel, whereas the Asn residues in the NPS/NSA motif in TbAQP2 form weak van der Waals interactions with the glycerol molecules, and Asn261 also makes a hydrogen bond to the backbone carbonyl of Leu129. The position of the intracellular loop between H2 and H3 is also conserved between TbAQP2 and PfAQP (Fig. 3c) with the backbone carbonyls of residues 126, 127 and 128 aligning and all forming hydrogen bonds in both channels to glycerol molecules. The conservation in structure is reflected in the conserved position of the glycerol molecules in the intracellular half of the channel in TbAQP2 and PfAQP, although the orientation of the hydroxyl groups of the glycerol molecules differs. A divergence in the position of the extracellular loop between H6 and H7 in TbAQP2 and PfAQP (Fig. 3c) leads to a wider channel in TbAQP2 and no ordered glycerol molecules are observed in the channel at this point, unlike in other aquaglyceroporins (Fig. 3d). Another difference between the transport of glycerol by TbAQP2 and PfAQP is that there is a water molecule coordinated between each glycerol molecule in PfAQP, whereas in TbAQP2 the glycerol molecules interact directly. Any role of water molecules in the transport of glycerol by TbAQP2 remains to be elucidated as the resolution of the current cryo-EM structure is insufficient for resolving them.

**Fig. 3.**
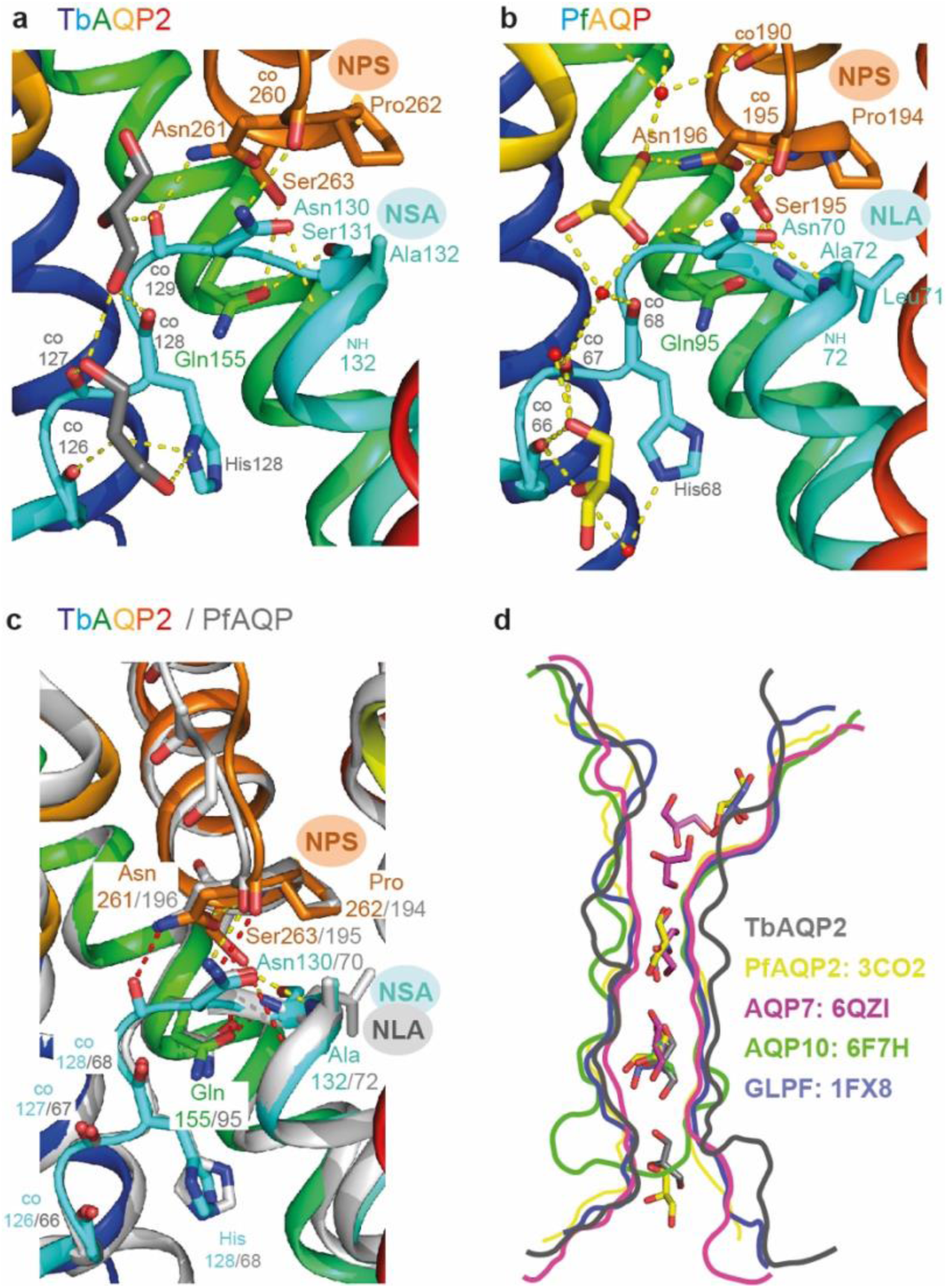
Comparison between conserved glycerol-binding regions of TbAQP2 and PfAQP. **a**, The structure of TbAQP2 is depicted (rainbow colouration) with residues shown that make hydrogen bonds (yellow dashed lines) to either glycerol (grey) or in the NPS/NSA motifs. CO, backbone carbonyl groups involved in hydrogen bond formation. **b**, The structure of PfAQP2 is depicted (rainbow colouration) with residues shown that make hydrogen bonds (yellow dashed lines) to either glycerol (yellow sticks) or water (red spheres), or in the NPS/NLA motifs. CO, backbone carbonyl groups involved in hydrogen bond formation. **c**, Structures of TbAQP2 (rainbow colouration) and PfAQP are superimposed to highlight structural conservation of the NPS and NSA/NLA motifs. **d**, Outline of channel cross-sections from aquaglyceroporins containing bound glycerol molecules (sticks): TbAQP2 (grey); PfAQP (yellow; PDB ID 3CO2); human AQP7 (magenta, PDB ID 6QZI); human AQP10 (green, PDB ID 6F7H); *Escherichia coli* GlpF (purple, PDB ID 1FX8).

### Molecular dynamics simulations show impaired pentamidine transport in mutants

To assess the dynamic properties of the structurally-resolved pentamidine binding pose, all-atom molecular dynamics (MD) simulations were performed on TbAQP2 in a tetrameric state (see Methods). In these simulations, both the protein and pentamidine are very stable (protein RMSD = 0.17 ± 0.01 nm and pentamidine RMSD = 0.16 ± 0.03 nm, see Fig. S9a,e) and pentamidine binding prevents water flux through the channel (Fig. 4a,b). The stability of the pentamidine is enabled by a number of extremely high occupancy interactions with specific residues lining the TbAQP2 pore (Fig. S9b). Hydrogen bond analysis reveals around 23 hydrogen bonds may be made between pentamidine and the channel (Fig. S10a,b). In contrast, water shows a different pattern of high occupancy interactions in the pore, with MD snapshots highlighting several key positions of where waters interact with the channel as they flow through (Fig. S10c). The presence of pentamidine both severely reduces the number of waters in the channel and slightly impacts their orientation (Fig. S10d,e). It was not possible to perform MD simulations of TbAQP2 in the presence of melarsoprol, because the presence of an arsenic atom causes force field parameterization issues.

**Fig. 4.**
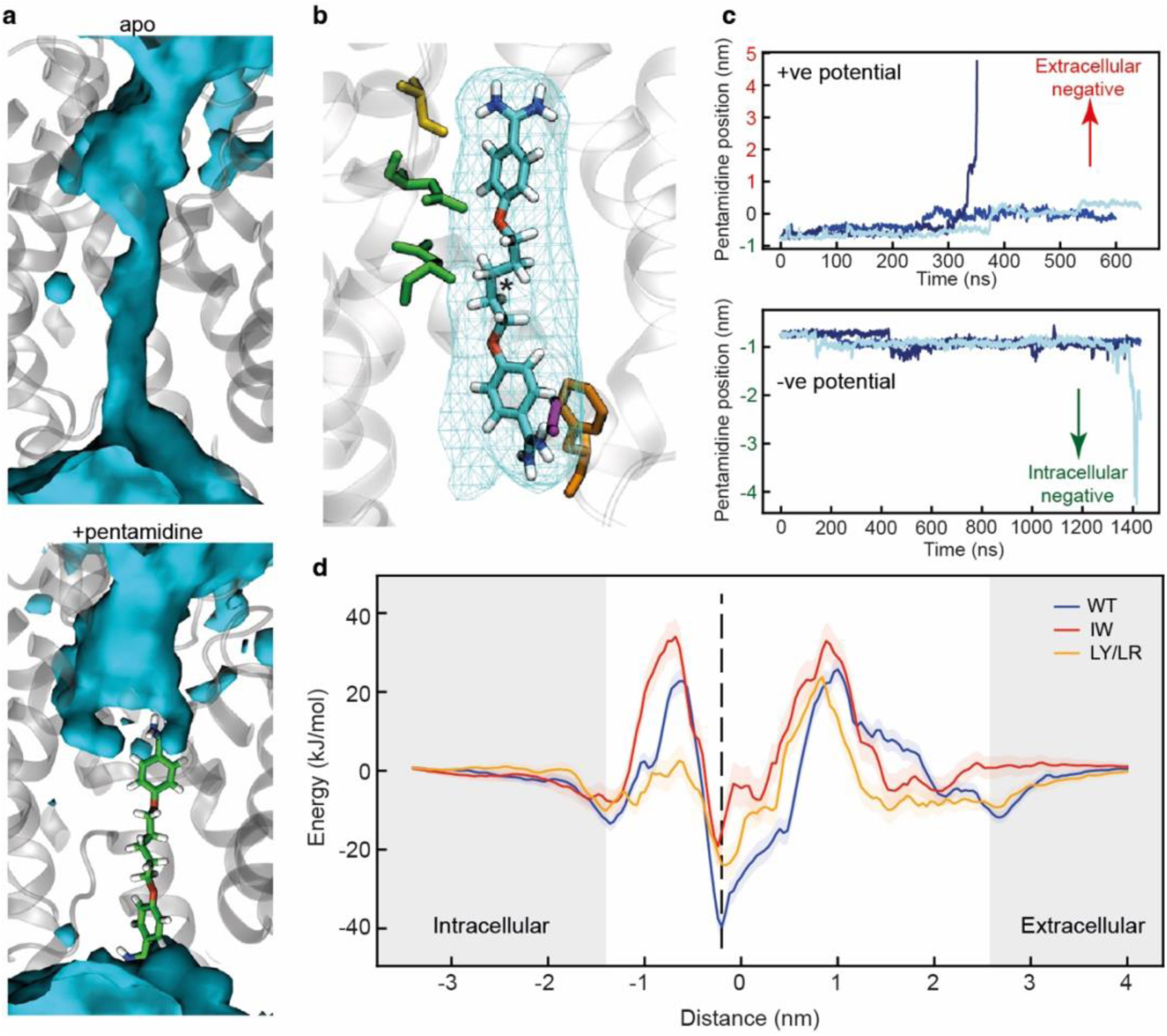
Molecular dynamics simulations of pentamidine-bound TbAQP2. **a**, Density of water molecules (blue) through the TbAQP2 central cavity from molecular dynamics. Bound pentamidine (sticks, green) abolishes the ability of TbAQP2 to transport water molecules. **b**, View of pentamidine (cyan, carbon; red, oxygen; blue, nitrogen; white, hydrogen) bound to TbAQP2. Cyan mesh shows the density of the molecule across the MD simulation. and the asterisk shows the position of the centre of mass (COM). **c**, Upon application of a membrane potential, the pentamidine position as defined by the COM moves along the z-dimension in relation to the COM of the channel, with three independent repeats shown in different shades of blue. The bottom graph is for the potential in the physiological direction (negative intracellular). **d**, Energy landscapes for pentamidine through the TbAQP2 central cavity as calculated using umbrella sampling. Separate calculations were made for monomeric WT TbAQP2 and TbAQP2 with I110W (IW) or L258Y/L264R (LY/LR) mutations. Each trace is built from 167 x 40 ns windows, with the histogram overlap and convergence plotted in Fig. S12a,b. The position of the membrane phosphates is shown as grey bars, and the structural binding pose is shown as a dotted line.

We then explored the translocation of pentamidine through the TbAQP2 channel. The positively charged state of pentamidine suggests that it might be translocated by the ΔΨ component of the plasma membrane potential. To test this, we applied a z-dimension electric field to simulations containing monomeric TbAQP2 in a membrane, using either a positive field (the opposite direction to the physiological membrane) or a physiologically-correct negative field polarity. In all simulations, the resulting potential causes the pentamidine to translocate through TbAQP2, and in some simulations (1 out of 3 repeats for each condition) the pentamidine fully leaves the channel (Fig. 4c). For the physiologically correct negative field, the pentamidine moves downwards (Fig. 4c), suggesting that the membrane potential is able to import pentamidine into the cell, consistent with experimental evidence that TbAQP2-mediated pentamidine uptake is sensitive to ionophores that depolarise the plasma membrane^10,17^.

### Drug resistant mutations in TbAQP2

*T. brucei* contains two related aquaglyceroporins, TbAQP2 and TbAQP3, with only TbAQP2 being able to transport pentamidine and melarsoprol into the trypanosome^11,12^. Drug resistance has arisen through recombination between the two genes and the expression of TbAQP2-AQP3 chimeras^13–15^. There are 68 amino acid residues that differ between TbAQP2 and TbAQP3 and the drug-resistant chimeras can contain 20-43 differences to TbAQP2 (Fig. S1). A number of these residues have been mutated individually in attempts to identify key residues involved in drug translocation and thus potentially resistance. At the intramembrane ends of the two transmembrane half-helices H3 and H7 are the aforementioned NSA/NPS motifs, Asn130-Ser131-Ala132 and Asn261-Pro262-Ser263, respectively (Fig. 3a). In aquaporins such as TbAQP3 these conserved residues are the canonical NPA/NPA motifs, and mutations were made (S131P+S263A) to convert the NSA/NPS motif in TbAQP2 to NPA/NPA. This resulted in a 95% reduction in pentamidine transport by the mutant, although this was still measurably higher than that observed in a TbAQP2 knockout cell line^17^ and a similar reduction in the transport of the melarsoprol analogue cymelarsan (Fig. S11). The cryo-EM structures show that both Ser131 and Ser263 are critical in stabilising this region of the transporter through specific hydrogen bonds with Gln155 and the backbone amine group of Ala132 (Fig. 3a) and do not make any direct interactions either to the drugs or the glycerol molecules. Thus, the decrease in pentamidine uptake observed in the S131P+S263A TbAQP2 mutant is probably more complex than previously thought, as it seems unlikely to have arisen through simple changes in transporter-drug interactions or a narrowing of the channel.

Based on a previous modelling study, several other mutations were predicted to potentially affect drug uptake in the TbAQP2-TbAQP3 chimeras^17^. Based on the cryo-EM structures, the mutated residues fall into three distinct groups: those not in the channel (I190T, W192G), those lining the channel and within van der Waals distance of pentamidine (I110W, L258Y, L264R) and those lining the channel, >3.9 Å from pentamidine and observed to make contacts to pentamidine during MD simulations (L84W/M, L118W/M, L218W/M). In all instances, there was a >90% decrease in pentamidine uptake) and in many cases no residual pentamidine transport was detected (e.g. I110W, L264R, L84W)^17^. To understand the molecular basis for the dramatic decreases observed in pentamidine transport, we performed molecular dynamics simulations on a subset of the mutants.

Three mutations were chosen for molecular dynamics simulations (I110W, L258Y, L264R), all of which were defective in uptake of cymelarsan (Fig. S11) as well as pentamidine^17^. These three residues all make high occupancy contacts to pentamidine in the MD simulations (Fig. S9b) and residues Ile110 and Leu264 make van der Waals contacts with pentamidine in the cryo-EM structure (Leu258 is 5.5 Å away from pentamidine). The double mutant L258Y+L264R was used in the MD simulations as this is found in a number of drug-resistant *T. brucei* strains such as P1000 and Lister 427MR^12^. The mutant I110W is found in the *T. brucei* drug resistant strain R25^28^, but in the absence of the L258Y+L264R double mutation (Fig S1). Therefore, we performed atomistic MD simulations of monomeric TbAQP2 with either the I110W (hereafter IW) or L258Y+L264R (hereafter LY/LR) mutations present, as well as with wild-type monomeric TbAQP2. Each protein was simulated both with pentamidine bound and removed (apo). The mutations do not affect TbAQP2 stability (Fig. S7e), but do destabilise pentamidine binding, causing it to shift away from the initial binding pose along the z-axis (Fig. S9c), suggesting a lowered affinity for the structural pose. The pore radii are unchanged by the mutations (Fig. S9d), meaning that this effect is likely to be a loss of specific interactions rather than a more general effect on protein structure.

We next computed energy landscapes for pentamidine moving along the z-axis through the channel using potential of mean force (PMF) calculations with umbrella sampling. The data for WT TbAQP2 reveals a clear energy well for the structurally resolved pentamidine at -0.2 nm (Fig. 4d and Fig. S11c). The depth of this well (-40 kJ/mol) suggests a very high affinity binding site, consistent with previous kinetic analyses^9,10,29^, and is much higher than previously investigated docked poses^17^ of pentamidine in models of TbAQP2. Notably, this energy well is much shallower for both the IW (-20 kJ/mol) and LY/LR mutants (-25 kJ/mol), representing a relative reduction in binding likelihood of about 2,800-fold and 400-fold, respectively. In biochemical terms, these changes would result in a Km shift from ∼50 nM to 50 or 300 µM, respectively, all but abolishing uptake at pharmacologically relevant concentrations; the experimental Km for pentamidine on wild type TbAQP2 is 36 nM^10^. The reduction in pentamidine binding likelihood is consistent with the increased z-axis dynamics seen in Fig. S9c. Flanking the central energy well are large energy barriers at about -0.7 nm and 1 nm (Fig. 4d), similar to those seen for urea translocation in HsAQP1^30^. These would presumably slow pentamidine entry into and through the channel, and previous studies did reveal a slow transport rate for pentamidine^10,29^. We note that previous studies using these approaches saw energy barriers of a similar size, and that these are reduced in the presence of a membrane voltage^17,31^. The energy barriers are highest in the IW mutant, and interestingly the inner (-0.7 nm) barrier is lower in the LY/LR mutant (Fig. S12e). In all cases, especially WT, small energy minima are observed at about -1.35 nm and +2.7 nm; these represent the drug binding at the entrance and exit of the channel (Fig. S12d). The differences in landscape between the WT and mutants helps explain how these mutations confer a resistance to pentamidine, mostly through reduced binding affinity.

## Discussion

It has only recently become accepted that the aquaglyceroporin AQP2 in *T. brucei* is acting as a *bona fide* transporter of pentamidine and melarsoprol that is essential for translocation of the drugs into the cell^17^. The cryo-EM structures of TbAQP2 identify the binding site for melarsoprol and pentamidine in a wide channel that has, at least for part of the trajectory, similar dimensions to channels in other aquaglyceroporins (Fig. 3d). The overall similarity between TbAQP2 and the structures of other aquaglyceroporins suggests a similarity in the mechanism of glycerol transport, despite differences in the amino acid residues lining the channel. It also suggests that there is scope for pharmacological targeting of other aquaglyceroporins, either for uptake of cytotoxic drugs or inhibition of glycerol uptake. Leishmaniasis, for instance, is another neglected tropical disease, caused by parasitic protozoa of the genus *Leishmania*; it is commonly treated by pentavalent antimonials meglumine antimoniate and sodium stibogluconate. The antimony that is released in the acidic parasitophorous vacuole enters the parasite through its AQP1 aquaporin^32^. Widespread antimonial drug resistance in *Leishmania donovani* in India has been linked to a frameshift mutation in LdAQP1^33^. In another example, *Plasmodium spp.* contain a single aquaglyceroporin and studies in mouse models have shown that a knockout of the aquaglyceroporin leads to a significant loss of virulence and reduction in erythrocyte infectivity by merozoites^34^, suggesting that a specific AQP inhibitor could be part of antimalarial treatments. Interestingly, pentamidine is a known antimalarial^35^ and although a substrate of the *P. falciparum* choline transporter, does not act through inhibition of that carrier^36^; inhibition/permeation of PfAQP by pentamidine has not been investigated to date. The structure determination of TbAQP2 highlights new opportunities for drug development targeting the aquaporin family to treat a number of parasitic diseases whose current treatments often suffer from severe side effects and growing problems in drug resistance. A thorough understanding of the interaction of extended, flexible drugs like pentamidine and melarsoprol with AQP channel linings could contribute to the design of new substrates able to penetrate not just protozoan but also human AQPs.

After this work was completed, a paper was published describing the structures of TbAQP2 bound to pentamidine and melarsoprol^31^. The structures of TbAQP2 are very similar to the ones described here (RMSD 0.6 Å), despite the environments of the purified channels being significantly different. In the work described here, TbAQP2 was purified in detergent and then incorporated into saposin A (SapA)^37^ lipid nanoparticles (see Methods), whereas Chen *et al.*^31^ incorporated TbAQP2 into lipid nanodiscs formed from a helical membrane scaffold protein (MSP)^38,39^. Lipid nanodiscs and nanoparticles are considered better mimetics of a lipid bilayer than detergent micelles, although biophysical studies indicate that there are significant differences between these lipidic structures and a biological membrane^40,41^. Comparison of the pentamidine in the different structures indicates good agreement in its position within the channel, although the pose of the drug does differ slightly (Fig. 5a). In contrast, there is a significant difference in the position and pose of melarsoprol between the different structures (Fig. 5b). Melarsoprol in the TbAQP2 structure from Chen *et al.*^31^ is shifted further towards the extracellular surface of the channel and is rotated approximately 180° about an axis in the membrane plane and centred on the arsenic atom (Fig 5b). This unexpected difference in orientation of melarsoprol in the two structures may perhaps not be so surprising as could be initially thought. The two structures represent the thermodynamically most stable states under the buffer conditions and detergent/lipid nanoparticles used for protein purification, which were different. In addition, the site of localisation of the drugs in the TbAQP2 channel is not a physiologically relevant ‘binding site’ as used to describe, for example, the orthosteric binding site in G protein coupled receptors where a hormone or neurotransmitter binds. Thus, evolution has not evolved a high-affinity binding site in the TbAQP2 channel and neither were the drugs designed to bind to TbAQP2. The channel merely facilitates the entry of the drugs into the cell through passive diffusion and it is not necessary for its mode of action to bind to TbAQP2 in a specific pose. Indeed, *a priori*, there is no reason to think that melarsoprol has to traverse the channel in any particular orientation, although under cellular conditions where a membrane potential is present it might be anticipated that the melamine moiety may enter the channel first given its net positive charge. Regrettably, in the absence of molecular dynamics simulations (due to the issues with parameterization of the arsenic atom) these ideas have to remain untested.

**Fig. 5.**
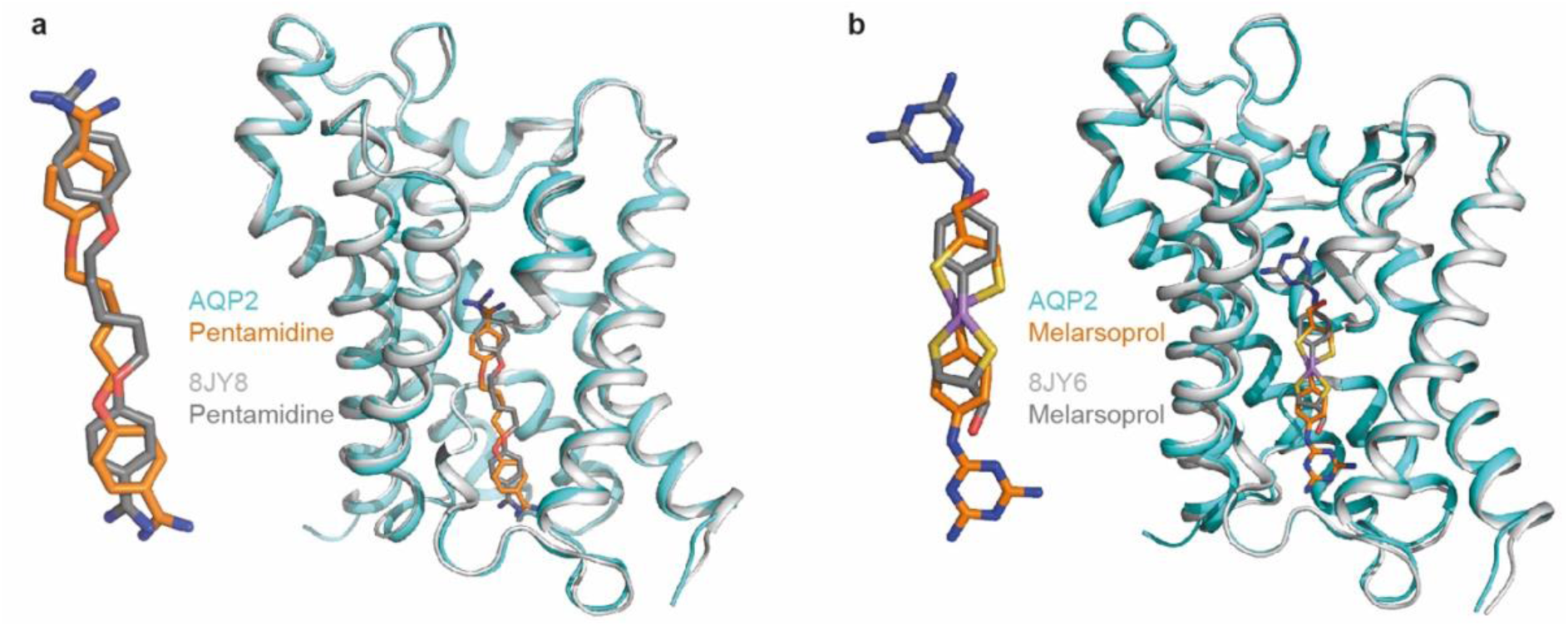
Comparison between drug-bound AQP2 structures. Superposition of an AQP2 protomer from this work (cyan) with a protomer determined by Chen *et al.*^31^ (grey), with the positions of drugs shown in stick representation: **a**, pentamidine; **b**, melarsoprol.

## Acknowledgements

S.N.W.’s lab was funded by a Sir Henry Dale fellowship from the Wellcome Trust and the Royal Society London (Grant Number 101234/Z/13/Z) and by the Isaac Newton Trust Cambridge (Grant Number 15.40(a)). The work in C.G.T.’s laboratory was supported by core funding from the Medical Research Council (MRC U105197215). T.S. was supported by an MRC-DTP PhD fellowship, a Cambridge Trust Vice-Chancellor’s Award, the Sackler fund and a King’s College Cambridge PhD overrun fund. R.A.C, M.S.P.S, and P.J.S are supported by Wellcome (208361/Z/17/Z). Aquaporin-related work in H.P.d.K. lab was supported by the Medical Research Council (Grant Number G0701258) and a BBSRC Impact Accelerator award (BB/S506734/1). R.W. was supported by the Wellcome Trust (208398/Z/17/Z). MC is a Wellcome Investigator (217138/Z/19/Z). This project made use of time on ARCHER2 granted via the UK High-End Computing Consortium for Biomolecular Simulation, HECBioSim (http://hecbiosim.ac.uk), supported by EPSRC (EP/R029407/1). In addition, simulations were performed using the computational facilities of the Advanced Computing Research Centre, University of Bristol (http://www.bristol.ac.uk/acrc/). This work benefited from access to the Instruct Image Processing Centre (I2PC), an Instruct-ERIC centre; financial support was provided by PID12540. For the purpose of open access, the MRC Laboratory of Molecular Biology has applied a CC BY public copyright licence to any Author Accepted Manuscript version arising.

## Author contributions

M.C. and S.N.W conceived the project. T.S. and S.N.W. cloned, overexpressed, purified, and analysed protein samples. J.N.B., P.C.-H., D.Y.C. and S.N.W. prepared grids, screened and collected data. M.G. and R.M. processed the raw data. M.M. and P.A. built the models and performed refinements. A.G performed further structural analyses. K.Y. and R.W. identified and fitted the substrates and performed refinements. J.M.C. advised on the processing and G.M. advised on the refinement process. V.S.C. performed hot spot analysis and T.L.B. advised. H.P.d.K. and M.C. advised the mutational studies. R.A.C. performed the molecular dynamics simulations, and R.A.C., P.S. and M.S. did the analysis of the MD simulation trajectories. M.A.U. performed cymelarsan uptake experiments. C.G.T. carried out structure analysis and S.N.W. managed the overall project. The manuscript was written by S.N.W., H.P.d.K., R.A.C. and C.G.T., and included contributions from all the authors.

## Data availability

Structures have been deposited in the Protein Data Bank (PDB; https://www.rcsb.org/), and the associated cryo-EM data have been deposited in the Electron Microscopy Data Bank (EMDB; https://www.ebi.ac.uk/pdbe/emdb/) and the Electron Microscopy Public Image Archive (EMPIAR; https://www.ebi.ac.uk/empiar/): SapA-TbAQP2-NP bound to glycerol (8OFZ, EMD-16864 and EMPIAR-11410); SapA-TbAQP2-NP bound to pentamidine (8OFY, EMD-16863 and EMPIAR-11412); SapA-TbAQP2-NP bound to melarsoprol (8OFX, EMD-16862 and EMPIAR-11411). There are no restrictions on data availability.

## METHODS

### Data reporting

Sample size was not predetermined by statistical methods, no randomization was applied to experiments and investigators were not blinded to experimental allocation or outcome assessment.

### Cloning and expression of wild-type AQP2

A construct encoding wild-type *T. brucei* AQP2 (residues 1–312) that included a tobacco etch virus (TEV) cleavage site, eGFP, and a decahistidine tag (TbAQP2-TEV-eGFP-His10) was generated and cloned into a pFastBac vector (Thermo Fisher) using standard ligation procedures. Recombinant pFastBac-TbAQP2-TEV-eGFP-His10 vector was transformed into Stellar Competent Cells (Takara) following the Bac-to-Bac Baculovirus Expression System manual (Thermo Fisher). Expression of AQP2 in *Spodoptera frugiperda* 9 (Sf9) cells (Thermo Fisher Scientific) was performed following the Bac-to-Bac Baculovirus Expression System manual (Thermo Fisher). The cells were then collected by centrifugation (10 min, 4°C, 3000 ×*g*), flash-frozen in liquid nitrogen, and stored at -80 °C until further use.

### Purification of AQP2 bound to either glycerol, pentamidine or melarsoprol and reconstitution into saposin-A nanoparticles (SapA-NP)

All steps described below were carried out at 4°C. *Sf*9 cells were defrosted on ice, resuspended in cold homogenization buffer (100 mM NaCl, 20 mM HEPES pH 7.5, 5% (v/v) glycerol, 2 mM PMSF, 1× Complete Protease Inhibitor Cocktail (Sigma)) and homogenised by applying 20 strokes in a Potter-Elvehjem glass homogenizer on ice. Membranes were solubilised using 1% (w/v) final concentration n-dodecyl-β-D-maltopyranoside (DDM, Anatrace) and a final concentration of 0.03% (w/v) cholesteryl hemisuccinate (CHS, Anatrace) for 4 hours at 4°C on a roller platform. Insoluble material was removed by ultracentrifugation (100,000 ×*g*, 60 min, 4 °C). The solubilisate was supplemented with 20 mM imidazole (final concentration) and incubated (16 h, 4°C) with 2 ml of Ni^2+^-NTA agarose resin (Qiagen) per litre of *Sf*9 cell culture. The resin was then applied to a gravity-flow column and washed three times with 20 column volumes washing buffer (100 mM NaCl, 20 mM HEPES pH 7.5, 2% (v/v) glycerol, 2 mM PMSF, 1× Complete Protease Inhibitor Cocktail (Sigma), 0.05% (w/v) DDM, 0.03% (w/v) CHS, 100 mM imidazole) and was eluted twice with one column volume elution buffer (100 mM NaCl, 20 mM HEPES pH 7.5, 2% (v/v) glycerol, 1× Complete Protease Inhibitor Cocktail (Sigma), 2 mM PMSF, 500 mM imidazole) each. Both eluates were combined and concentrated in an Amicon Ultra 4 mL centrifugal filter (100,000 MWCO, Merck) and aggregates were removed using a syringe-driven filter units (Millex-GV, 0.22 μm, PVDF, 13 mm, Merck). The filtrate was loaded at 0.4 ml/min onto a Superdex Increase S200 10/300 (GE Healthcare) size exclusion chromatography (SEC) column, pre-equilibrated with 0.05% (w/v) DDM, 100 mM NaCl, 20 mM HEPES pH 7.5. The peak fractions containing AQP2 were pooled and concentrated as above. Protein concentration was estimated by the Pierce BCS assay according to the manufacturer’s instructions.

SapA-NPs were prepared and stored in SapA-NP buffer consisting of 150 mM NaCl, 50 mM sodium acetate pH 4.8)^42^. Purified TbAQP2 tetramer was combined at a TbAQP2:SapA:lipid molar ratio of 0.1:1:10 and dialysed (20 h, 4°C) against SapA-reconstitution buffer (100 mM NaCl, 20 mM HEPES pH 7.5). SapA-NPs, unincorporated TbAQP2 and SapA were removed by SEC and SapA-TbAQP2-NP tetramer complex was concentrated using Amicon Ultra centrifugal filters (100,000 MWCO; Merck). Melarsoprol or pentamidine was added to the purified SapA-TbAQP2-NP tetramer complex (final concentration 1 mM, 2 h, 4°C) applied to a Superdex Increase S200 10/300 SEC column (GE Healthcare) and fractions containing the complex were concentrated using Amicon Ultra centrifugal filters (100,000 MWCO; Merck) to final concentrations between 0.8 mg/ml and 1.6 mg/ml. Final samples were centrifuged (1 h, 18,000 ×*g*, 4°C) immediately prior to cryo-EM grid preparation.

### Vitrified sample preparation and data collection

Cryo-EM grids were prepared by applying 2 μl of purified SapA-TbAQP2-NP with either glycerol, pentamidine or melarsoprol bound (0.8-1.6 mg/ml) onto either glow-discharged PEGylated UltrAuFoil R1.2/1.3 300 Mesh Gold grids (Agar scientific, No. AGS187-3) or glow-discharged Quantifoil R1.2/1.3 300 Mesh Gold grids (Agar scientific, No. AGS143-8). Grids were blotted with filter paper for 4 s before plunge-freezing in liquid ethane (at −181 °C) using an Thermo Fisher Scientific Vitrobot Mark IV (95% relative humidity, 4°C). All cryo-EM datasets were collected on Titan Krios microscopes (Thermo Fisher Scientific) operating at 300 kV. SapA-TbAQP2-NP with glycerol was collected at the Nanoscience Centre (University of Cambridge) with a Falcon 3 detector to obtain dose-fractioned movies. 2881 movies were recorded with 75,000× magnification at 1.07 Å per pixel using an exposure time of 60 s resulting in a total exposure of (30 e/pixel) and a target defocus range of -1.4 and -3.2 μm. Datasets of SapA-TbAQP2-NP with bound pentamidine or melarsoprol were collected at the cryo-EM facility at the Department of Biochemistry, University of Cambridge; images were recorded using a K2 detector in counting mode, operating with GIF Quantum LS imaging filter (Gatan) with a slit width of 20 eV to obtain dose-fractioned movies with a 100 µm objective aperture. 2400 movies were recorded for SapA-TbAQP2-NP with pentamidine bound and 3000 movies for melarsoprol bound complex. Micrographs for both datasets were recorded at a magnification of 130,000× (1.07 Å per pixel) as dose-fractioned movie frames with an exposure time of 15 s and 12 s, respectively, resulting in a total exposure of 54 e/Å^2^ and 57.5 e/Å^2^ with target defocus ranges of -1.5 and -3.3 μm, and -1.2 and -3 μm, respectively.

### Cryo-EM data processing and 3D reconstruction

Cryo-EM data processing was conducted inside the Scipion platform^43^. Image stacks (2,881 glycerol-bound, 4,086 melarsoprol-bound and 3,262 pentamidine-bound) were subjected to beam-induced motion correction using Relion’s implementation of MotionCor2^44^ by dividing each frame into 5 × 5 patches. CTF parameters were estimated from dose-weighted micrographs as follows: estimations made with GCTF^45^ with equiphase averaging and with CTFFIND-4.1^46,47^ were compared using «xmipp-ctf consensus». Outputs from these consensus (micrographs for which the two algorithms agreed in their estimations) undergone an additional round of ctf estimation with «xmipp-ctf estimation»^48^, followed by a second round of xmipp-ctf consensus for micrograph curation based on defocus, astigmatism, resolution and ice thickness. As a result, the number of micrographs for downstream analysis was 2102, 3586 and 2983 for SapA-TbAQP2-NP bound to either glycerol, melarsoprol or pentamidine, respectively.

Autopicking was performed using crYOLO^49^, that yielded 849,607 particles, 1,927,102 particles and 1,263,534 particles for SapA-TbAQP2-NP bound to either glycerol, melarsoprol or pentamidine, respectively. Particles were extracted in a box-size of 250 px at the original sampling rate of the micrographs (1.07 Å/px) and subjected to several rounds of 2D classifications with cryoSPARC^50^. To further curate the particle set (ligand-free: 100,075; melarsoprol: 202,752; pentamidine: 114,206). Particles were re-extracted with a box of 320 px before 3D classifications.

For the glycerol-bound dataset, an *ab initio* 3D model was generated using stochastic gradient descent algorithm implemented in cryoSPARC, followed by non-uniform refinement^51^ with C4 symmetry and a static mask encompassing the transmembrane domains and nanodiscs belt, while excluding the mobile GFP tags. With these settings, a 3.2 Å-resolution map was obtained. For the melarsoprol and pentamidine datasets, six *ab initio* models were generated. Particles from the best classes (melarsoprol: 126,551; pentamidine: 83845) were used for non-uniform-refinement with the same strategy as for the glycerol-bound particles, yielding a 3.2 and 3.7 Å-resolution map, respectively.

The global resolution was calculated following the FSC gold standard threshold of 0.143. The refined maps were sharpened using deepEMhancer^52^ to facilitate model building. Finally, voxel size for melarsoprol and pentamidine was re-adjusted to 1.042 Å/px using Relion post-process^53^. Local resolution was estimated with MonoRes^54^.

### Structure determination and model refinement

The monomer structure of TbAQP2 was predicted using the AlphaFold2 program as a starting point for further refinement^55^. The final cryo-EM map was converted to an mtz file with a range of sharpening and blurring options using CCPEM^56,57^. EMDA was used for fitting maps^58^. Real-space refinement of the atomic coordinates into the cryo-EM map was performed using Phenix^59^ and Coot software^60^. Finally, models in the asymmetric unit were refined against unsharpened and unweighted half maps using Servalcat Refmac5 pipeline^61^ with *C*4 symmetry constraints. The reference structure restraints were prepared with ProSmart^62^ using AlphaFold2 predicted models from the AlphaFold DB^63^. The final models were evaluated for geometry, close contacts, and bond parameters using MolProbity^64^. Graphic representations of the fitted coordinates into electron density and the final cryo-EM map were generated using PyMOL^65^ and UCSF ChimeraX^66,67^ packages.

### Molecular dynamics simulations

The model for MD simulations was based on the structure of TbAQP2 with pentamidine bound and was built into simulation boxes using CHARMM-GUI^68,69^ (see Table S2 for a list of systems built). Protein atoms were described with the CHARMM36m force field^70,71^ TbAQP2 was built either as a tetramer or a monomer. Side chain pKas were assessed using propKa3.1 and side chain side charge states were set to their default^72^. The pentamidine molecule used existing parameters available in the CHARMM36 database under the name PNTM with a charge state of +2 to reflect the predicted pKas of >10 for these groups^73^ and in line with previous MD studies^17^. For the ligand-free simulations, the pentamidine molecule was removed prior to simulation.

The proteins were built into membranes comprising 20% cholesterol and 80% POPC, solvated with explicit water molecules using the TIP3P model, and neutralised with K^+^ and Cl^-^ to 150 mM. Box sizes were initially set to 10 × 10 × 10 nm for the tetrameric systems and 6.5 × 6.5 × 10 nm for the monomeric systems. The use of an all-atom fixed charge force field with explicit solvent and membrane is optimal to understanding the protein-drug interactions central to this study. Each system was minimized and equilibrated according to the standard CHARMM-GUI protocol^68,69^. Production simulations using random seeds were run in the NPT ensemble with 2 fs time steps, temperatures held at 303.5 K using a velocity-rescale thermostat and a coupling constant of 1 ps, and pressure maintained at 1 bar using a semi-isotropic Parrinello-Rahman pressure coupling with a coupling constant of 5 ps^74,75^. Short range van der Waals and electrostatics were cut off at 1.2 nm, and the particle mesh Ewald (PME) method was used for long-range Lennard-Jones interactions^76^. Hydrogen bonds were constrained using the LINCS algorithm.

Membrane potential simulations were run using the computational electrophysiology protocol. An electric field of 0.1 V/nm was applied in the z-axis dimension only, to create a membrane potential of about 1 V (see Fig. S12a). Note that this is higher than the physiological value of 87.1 ± 2.1 mV at pH 7.3 in bloodstream *T. brucei*^77^ and was chosen to improve the sampling efficiency of the simulations. The protein and lipid molecules were visually confirmed to be unaffected by this voltage, which we quantify using RMSF analysis on pentamidine-contacting residues (Fig. S12b).

Potential of mean force (PMF) calculations were built from initial steered MD pulls using an umbrella potential along a coordinate of the COM of the pentamidine molecule and the COM of select residues in TbAQP2 which form the pentamidine binding site (Leu84, Ile110, Val114, Leu122, Gly127, Leu129, Leu218, Val222, Phe226, Leu244, Val245, Ala259, Met260, and Leu264). Pulls were run in the z-axis in either a positive or negative direction, at a rate of 1 nm per ns and with a force constant of 1000 kJ/mol/nm^2^. Snapshots were taken along the coordinates every 0.05 nm, using a script from Dr. Owen Vickery (https://doi.org/10.5281/zenodo.3592318), with a total of 167 snapshots to construct the reaction coordinate. End points for the coordinate were determined based on the pentamidine no longer making any contacts (<0.4 nm) with any atoms from the protein or membrane. Simulations were run for 5 ns per window with a 5,000 kJ/mol/nm^2^ umbrella restraint imposed to keep the system in place with respect to the reaction coordinate. An atom in Leu218 was chose as a reference for the treatment of periodic boundary conditions. Forces were written every 1 ps. PMF profiles were constructed using the weighted histogram analysis method (WHAM)^78^ implemented in *gmx wham*^79^ between -3.5 nm to 4.5 nm of the coordinate and with cyclisation applied. The first 50 ns of each window was discarded as equilibration time and 200 Bayesian bootstraps were used to estimate the PMF error. Adequate coverage of the landscape was assessed based on histogram overlap (Fig. S10c), and convergence was determined based on re-running the WHAM calculations on longer windows until the landscapes converged (Fig. S10d).

All simulations were run in Gromacs 2020.3 (https://doi.org/10.5281/zenodo.7323409)^80^. Data were analysed using Gromacs tools and VMD^81^. Pentamidine interactions with TbAQP2 were assessed using PyLipID^82^. Pore radii were assessed using the CHAP package^83^. Plots were made using Matplotlib^84^ or Prism 9 (GraphPad). MD input and output coordinate files are available at https://osf.io/zc235/.

### Docking and hotspots experiments

Fragment Hotspot Maps (FHMs) is a powerful tool for predicting the binding hotspots of small molecules on protein surfaces^21^. FHMs are based on the concept that certain chemical fragments are more likely to interact with specific regions of a protein’s surface, or ‘hotspots’, than others. By systematically mapping the interactions between a library of fragment molecules and a target protein, FHMs can identify key hotspots and help guide the design of new small molecules with improved binding affinity and specificity. The development of FHMs has been driven by advances in computational methods for predicting protein-ligand interactions, as well as by improvements in experimental techniques for mapping the 3D structure of protein-ligand complexes. FHMs have been used to guide drug discovery efforts in a variety of therapeutic areas, including oncology, infectious diseases, and neurology.

One key advantage of FHMs is that they can help to identify "druggable" regions of a protein’s surface that may be otherwise difficult to target with small molecules. For example, many protein-protein interaction interfaces are large and flat, making it challenging to design small molecules that can disrupt these interactions. FHMs can help to identify the specific regions of the interface that are most important for binding and guide the design of molecules that can effectively target these regions. Another advantage of FHMs is that they can be used to predict the binding modes of small molecules that have not yet been synthesized or tested experimentally. This can be particularly useful in early-stage drug discovery, where computational methods are often used to screen large databases of compounds for potential hits. Overall, FHMs represent a powerful approach for predicting and optimizing small molecule binding to protein surfaces.

We used FHMs program to generate hotspots for AQP2 monomer and probe protein’s potential ligand binding sites. The superposition of the final structures and the predicted hotspot maps show a clear overlap between the hotspots and the two drugs (Fig. S2).

### Lysis assay with arsenical drugs

The effects of cymelarsan and phenylarsine oxide (PAO) on *T. brucei* Lister 427 bloodstream forms was measured as the decrease in cell absorbance at λ 750 nm over time, based on the reduction of cell motility and increased cell lysis leading to reduced light scatter and absorbance in the cuvette^85^, exactly as described^86^. Briefly, 2×10^6^ cells in 200 µL buffer was placed in the wells of a 96-well plate and absorbance recorded in a PHERAstar plate reader (BMG Labtech, Durham, NC, USA) for 10 min before the addition of 20 µL of 100 µM of either cymelarsan or PAO in aqueous buffer, or buffer alone.

## Supplementary Table and Figure legends

**Fig. S1.**
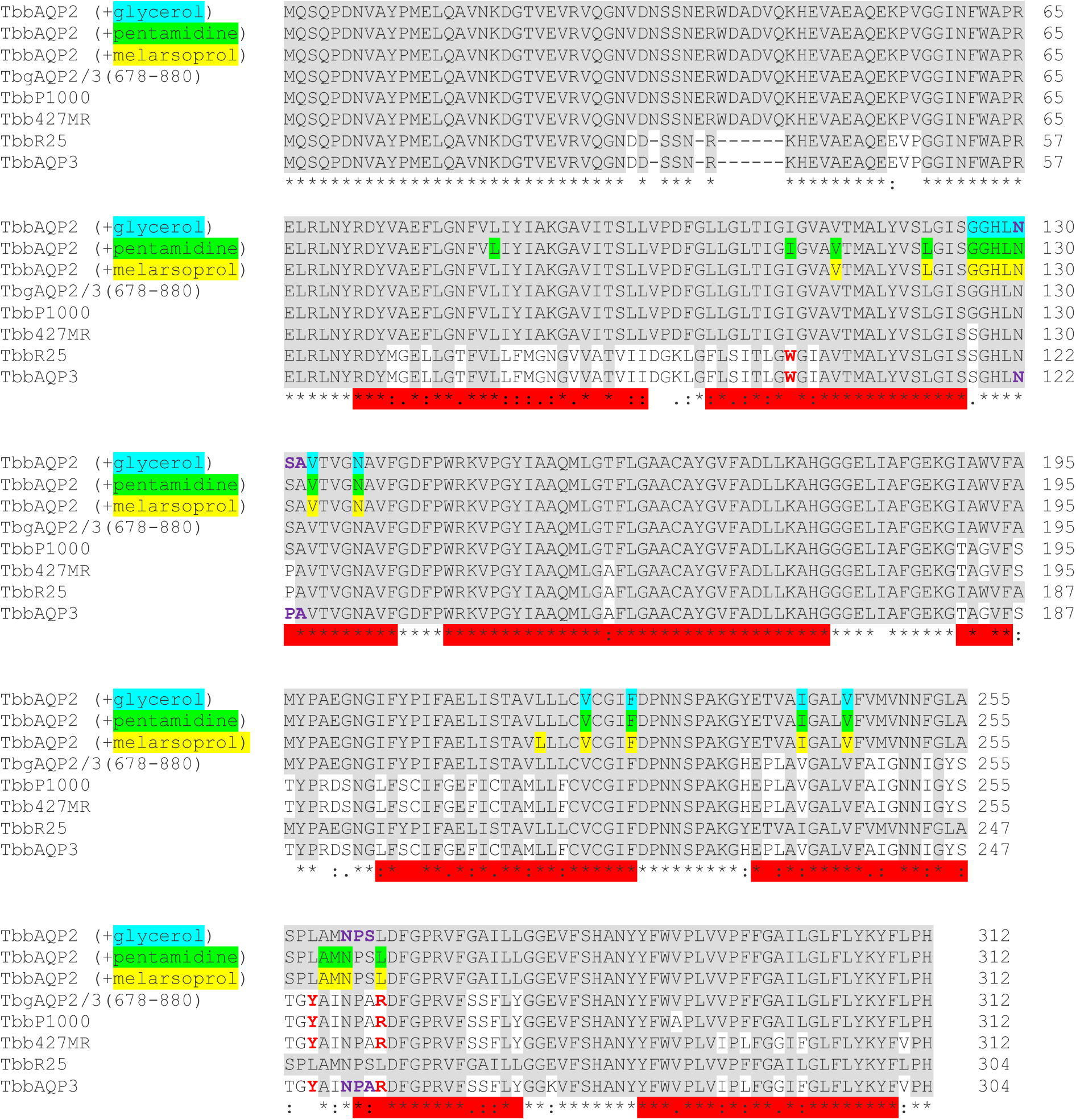
Alignment of aquaporin amino acid sequences. Amino acid sequences of AQP2 and AQP3 from *Trypanosoma brucei brucei* (Tbb) are aligned with chimeric aquaporins from drug resistant strains (P1000, 427MR, R25) and a chimera from drug-resistant *T. b. gambiense* (TbgAQP2/3). Three identical sequences are shown of TbbAQP2 with residues making contact to either glycerol, pentamidine or melarsoprol highlighted (cyan, green or yellow, respectively). Red bars represent transmembrane regions. Residues highlighted in grey are conserved in all sequences. The conserved NSA/NPS motif in TbAQP2 and the analogous NPA/NPA motif in TbAQP3 are in purple font and the residues mutated for the MD simulation studies (I110W, L258Y and L264R) are in red. Sequence information was obtained from Munday et al.^12^ except R25^28^ and TbgAQP2/3 (678-800)^15^.

**Fig. S2.**
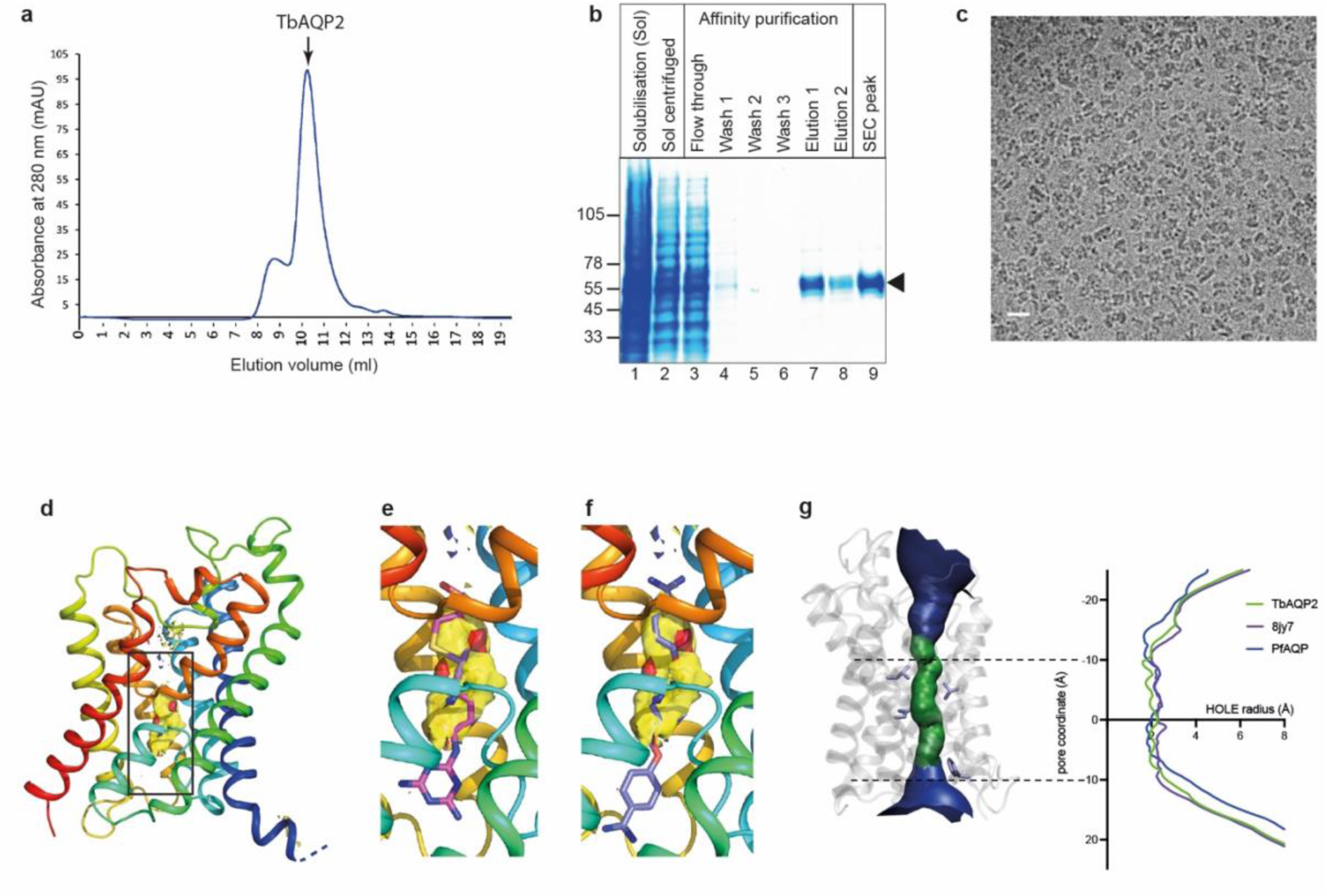
Purification of TbAQP2 and identification of the drug binding site by hotspot analysis. **a**, SEC trace of affinity purified TbAQP2. **b**, Coomassie-stained SDS-PAGE gel of fractions during detergent solubilisation (Sol), Ni^2+^-affinity chromatography and from the SEC column. **c**, Cryo-EM electron micrograph showing the particle distribution of TbAQP2. **d**, A region of TbAQP2 was identified as being most likely to bind a ligand is shown (yellow surface) is mapped on to the structure of TbAQP2 determined here (rainbow colouration. **e**, as in **d**, but also containing pentamidine (cyan sticks). **f**, HOLE^87^ trace for our resolved TbAQP2 structure, compared to a previously released TbAQP2 structure (‘8yj7’)^31^ and *Plasmodium falciparum* AQP^24^.

**Fig. S3.**
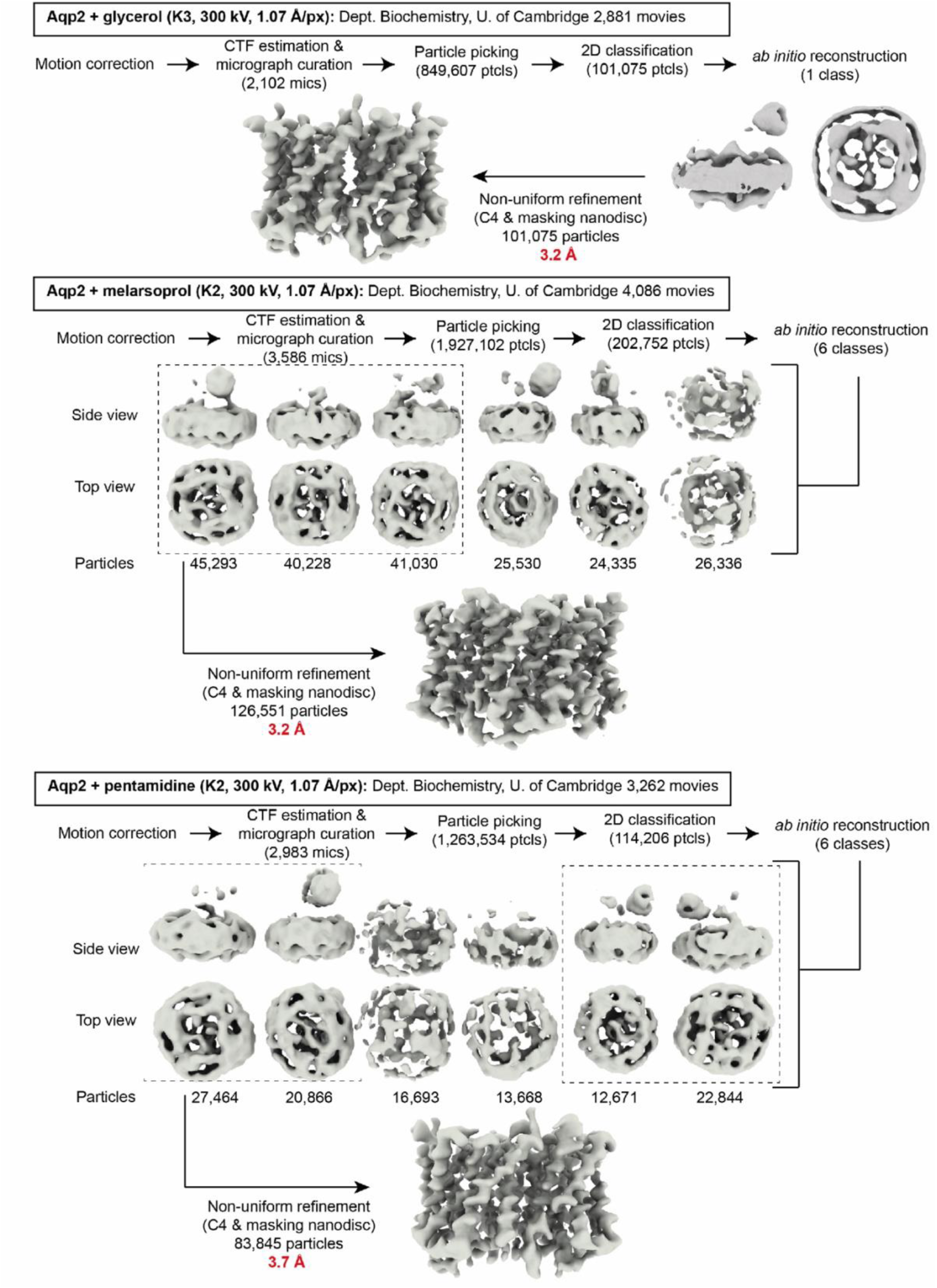
Flow chart of the cryo-EM image processing and structure determination. See the Methods section for the computer programmes used for each step and for further details.

**Fig. S4.**
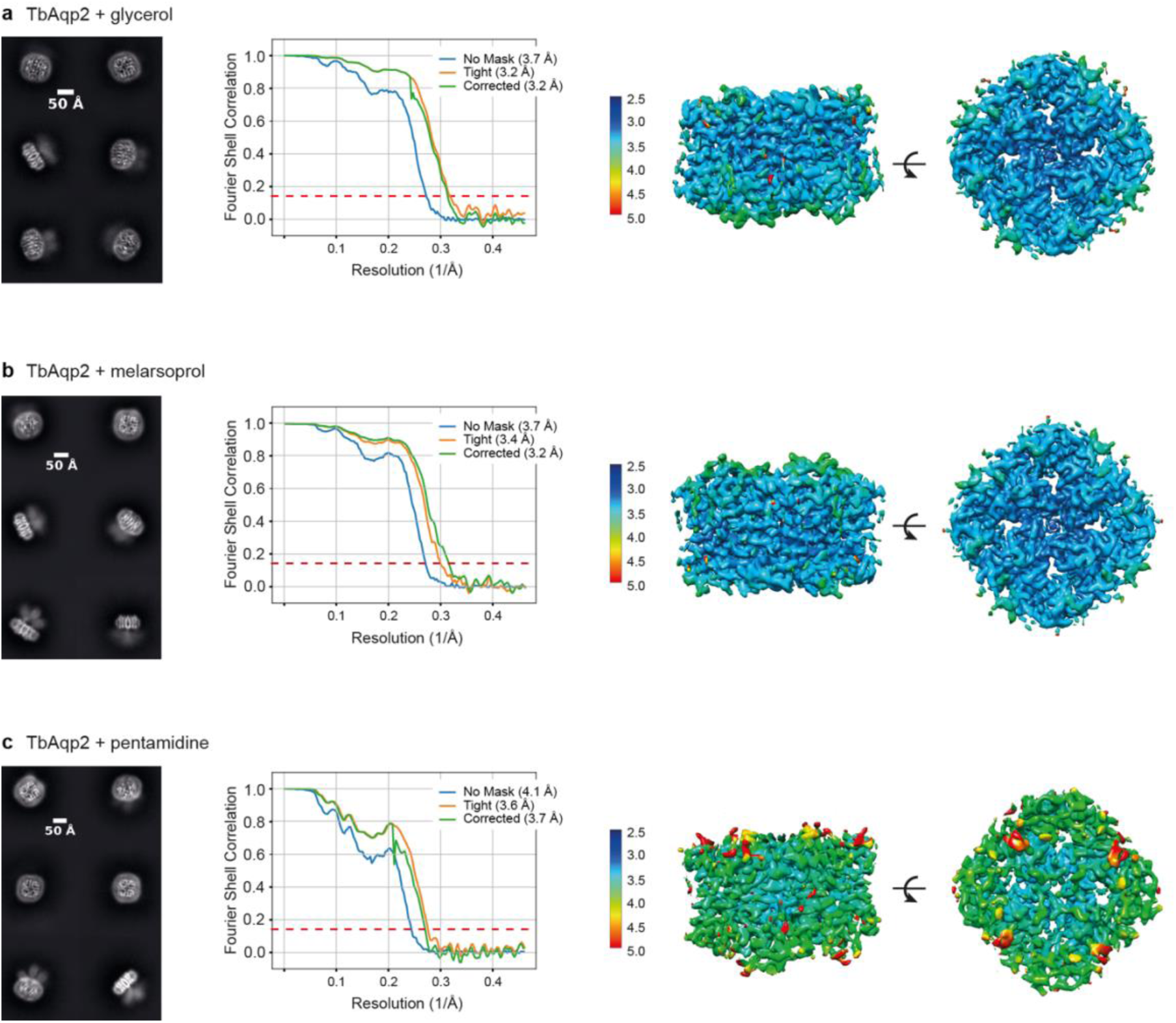
Local resolution maps of TbAQP2 structures. **a**-**c**, 2D class averages for each of the structures (left panels), FSC curves of the reconstructions with estimates for resolution determined using an FSC of 0.143 as implemented in cryoSPARC (middle panels) and local resolution estimations as calculated by MonoRes (right panels).

**Fig. S5.**
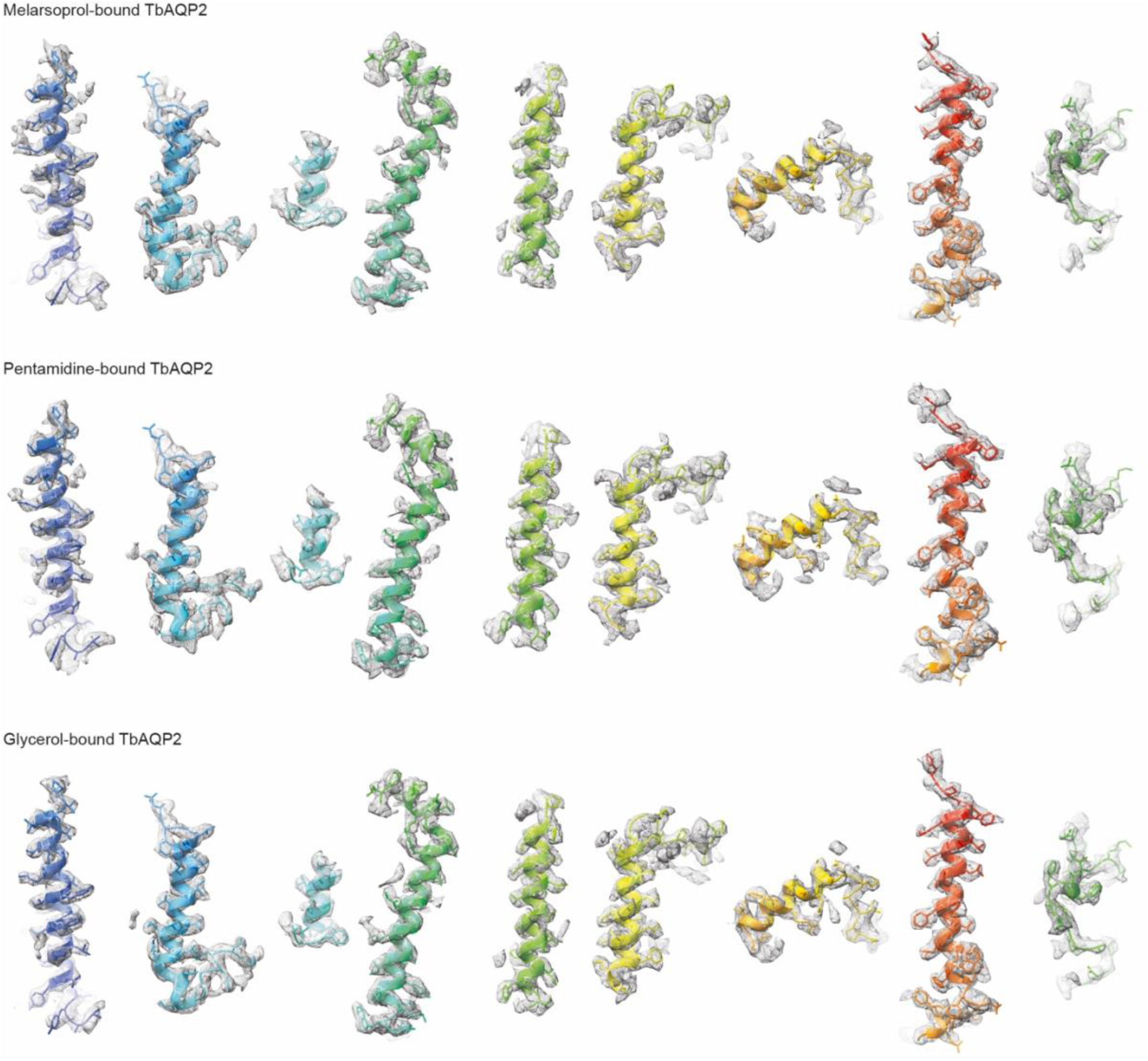
Density maps of secondary structure elements in the TbAQP2 structures. Colouring is the same as in Fig 1b.

**Fig. S6.**
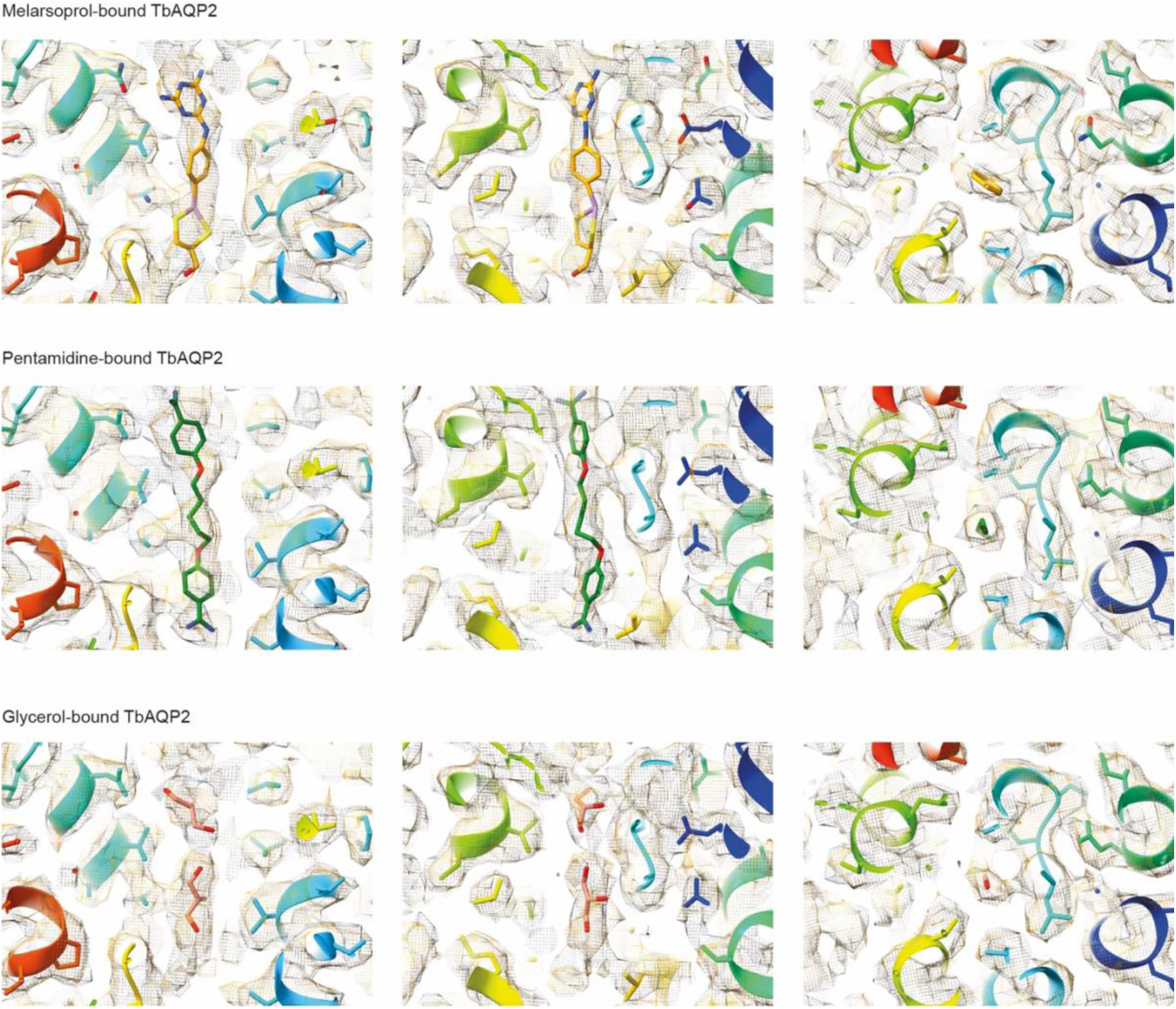
**Density half maps of ligands bound in the TbAQP2 structures.**

**Fig. S7.**
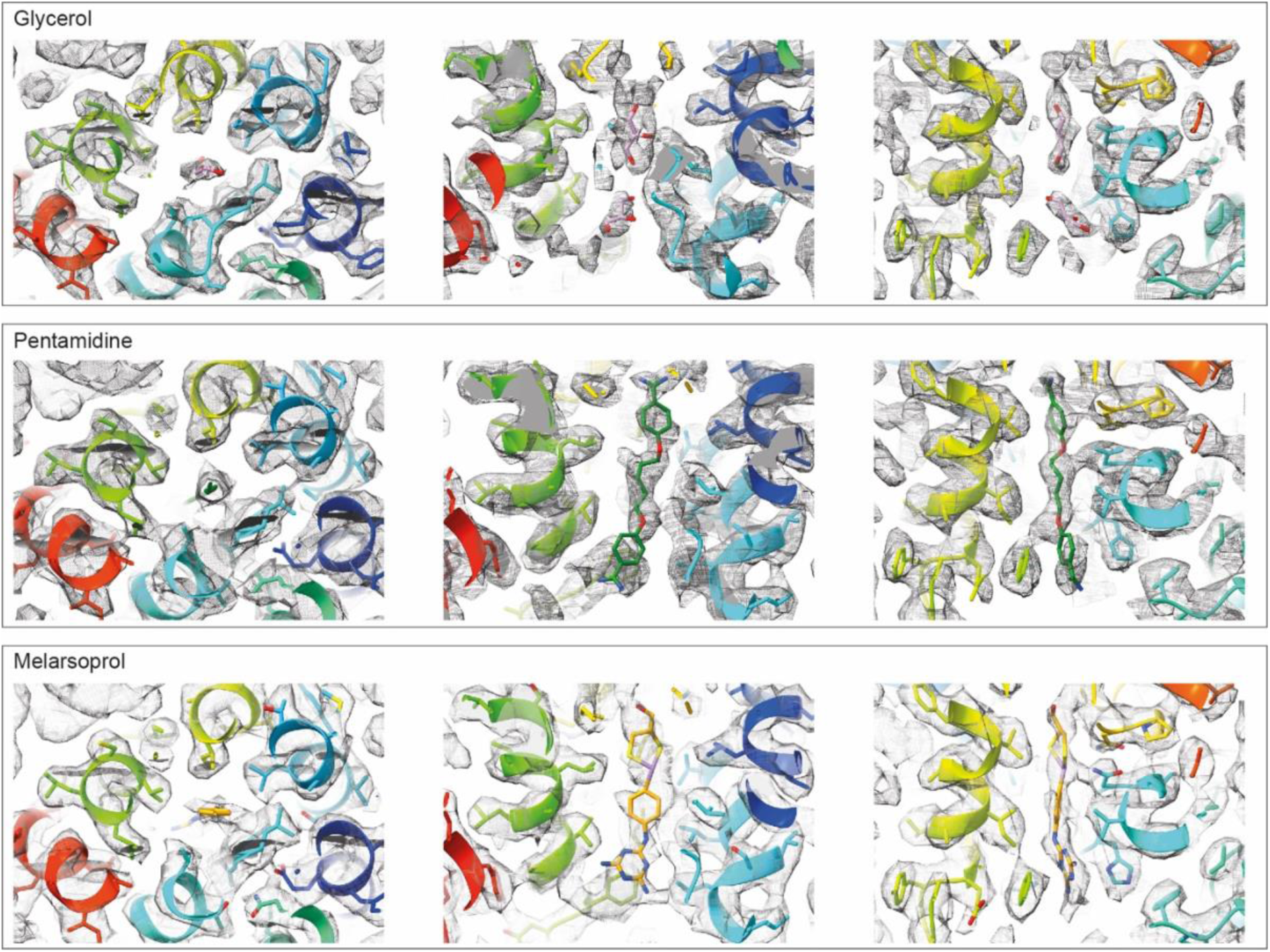
**Depictions of the side chain and substrate densities.**

**Fig. S8.**
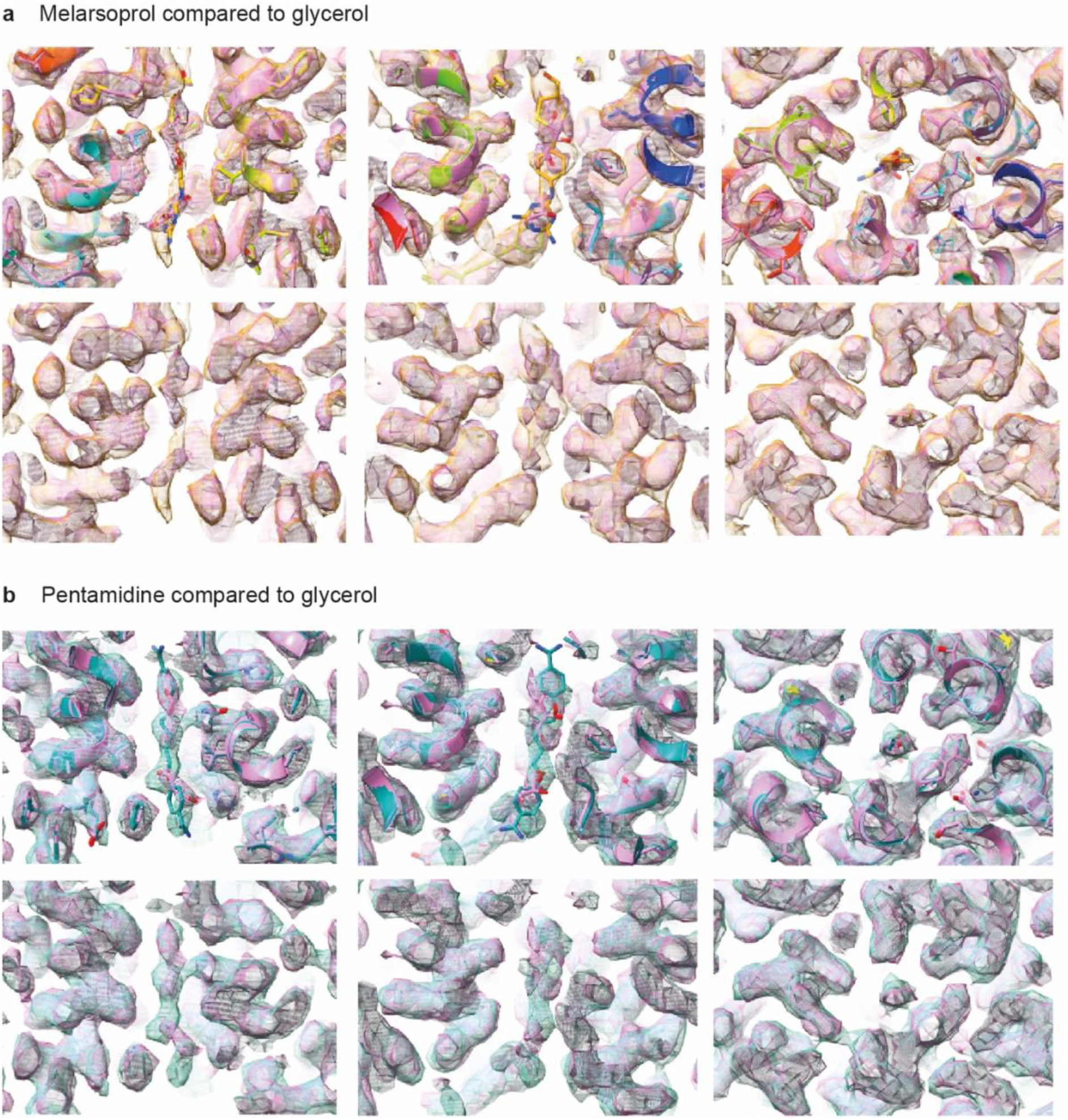
Comparisons between substrate densities. **a**, Comparison between melarsoprol and glycerol. **b**, comparison between pentamidine and glycerol.

**Fig. S9.**
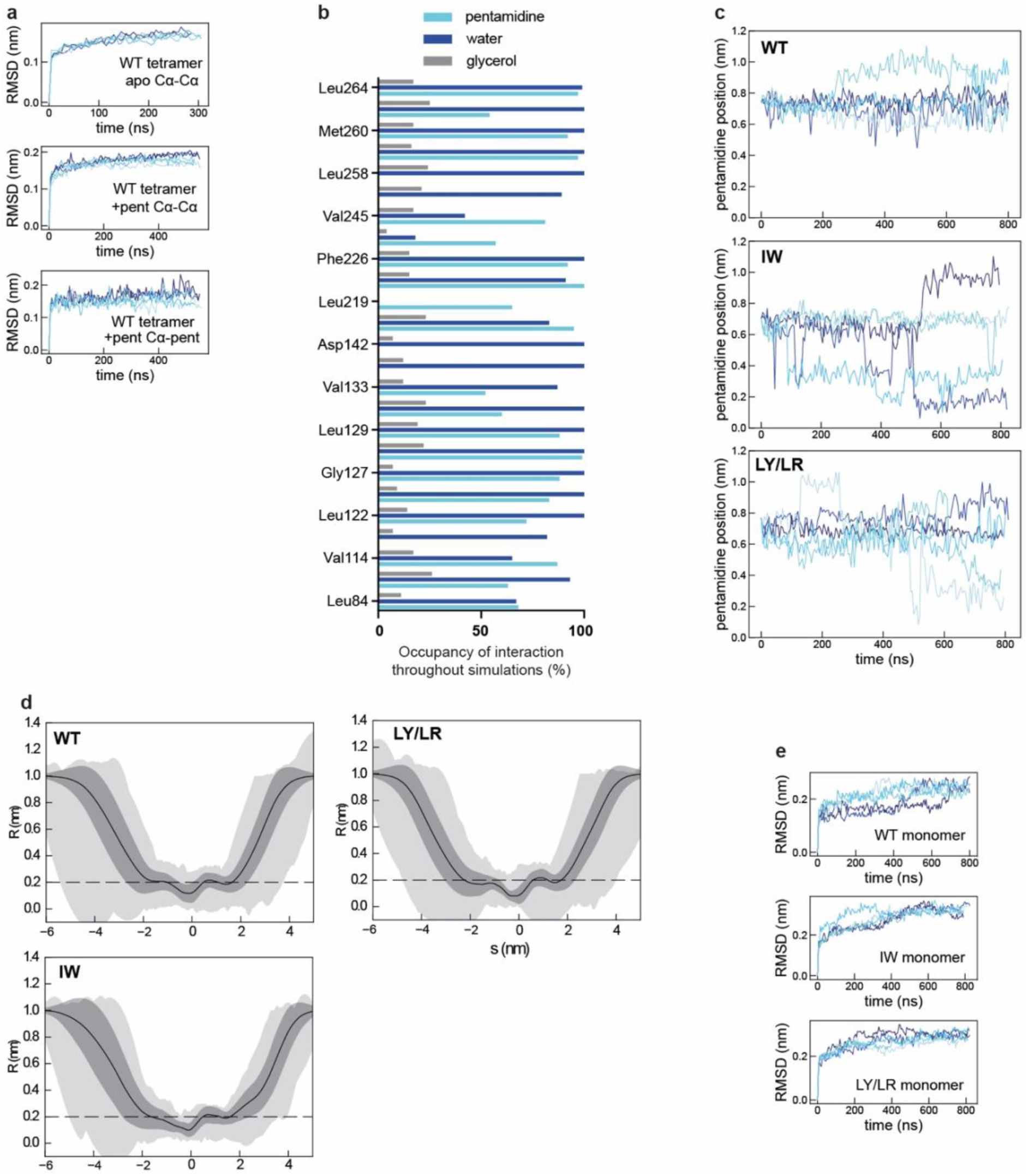
Molecular dynamics simulations. **a**, RMSDs for simulations of tetrameric WT TbAQP2. Trajectories were fitted on the Cα atoms and RMSDs were calculated for either Cα atoms for apo TbAQP2 (top), Cα atoms for pentamidine (pent) bound TbAQP2 (middle), or for the pentamidine molecule itself, i.e. in relation to the Cα of the channel (bottom). Five independent repeats are shown as blue traces. **b**, Percentage of time (occupancy) a given residue is within 3.9 Å of either pentamidine, water or glycerol during the MD simulations. **c**, Plotting the z-axis position of pentamidine when bound to monomeric TbAQP2 in a WT, IW, or LY/LR background. **d**, Pore radius profiles from simulations of apo monomeric WT, IW, or LY/LR TbAQP2. Analyses were run using the CHAP package^83^ with traces showing the mean (black line), standard deviation (dark grey), and range (light grey) for all frames over 5 x 800 ns of simulation. **e**, As panel **a**, but for Cα atoms from monomeric TbAQP2 in a WT background, or with I110W (IW) or L258Y/L264R (LY/LR) mutations.

**Fig. S10.**
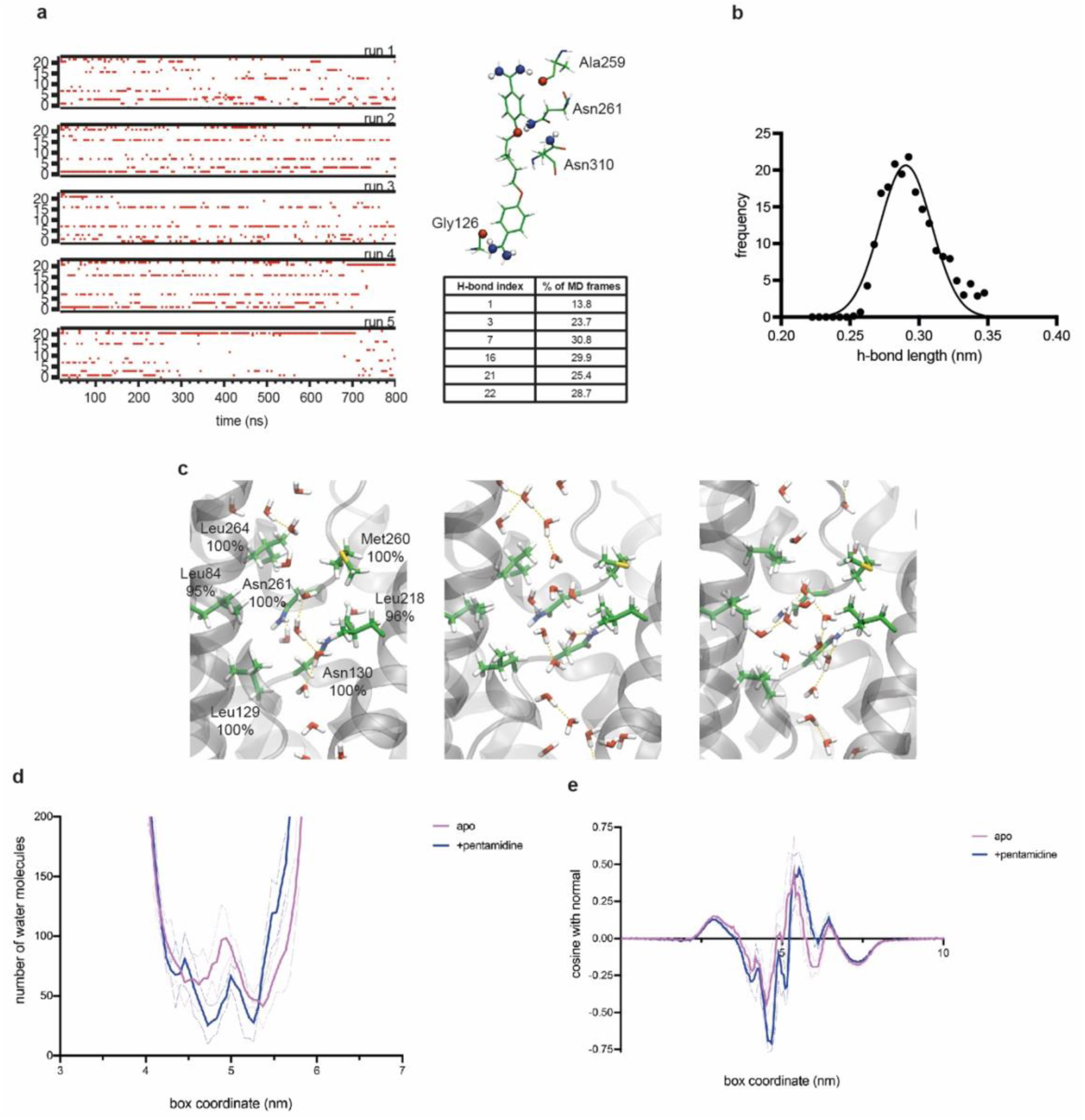
Water in molecular dynamics simulations. **a**, hydrogen-bond analysis from *gmx hbond* between the pentamidine and channel. Across the 5 independent MD runs of the monomeric TbAQP2 MD simulations, 23 hydrogen-bonds are formed, but only 6 appear to be high occupancy. These are highlighted on the image along with their index number, and their quantification is shown below. **b**, hydrogen-bond distances from the bonds in panel **a** plotted as a frequency plot. The average bond length was 0.29±0.02 nm, suggesting moderate strength. **c**, snapshots from MD showing water progression through the TbAQP2 channel in the absence of pentamidine. Three snapshots are shown. Residues that form especially high contact with water molecules (values given in brackets) are highlighted in each snapshot. Water-water hydrogen bonds, as computed using VMD, are shown in yellow dashed lines. **d**, number of water molecules along the TbAQP2 channel ±pentamidine, as computed using *gmx h2order* on the monomeric TbAQP2 MD simulations. **e**, orientation of the waters in each of the slices along with channel, as computed using *gmx h2order*.

**Fig. S11.**
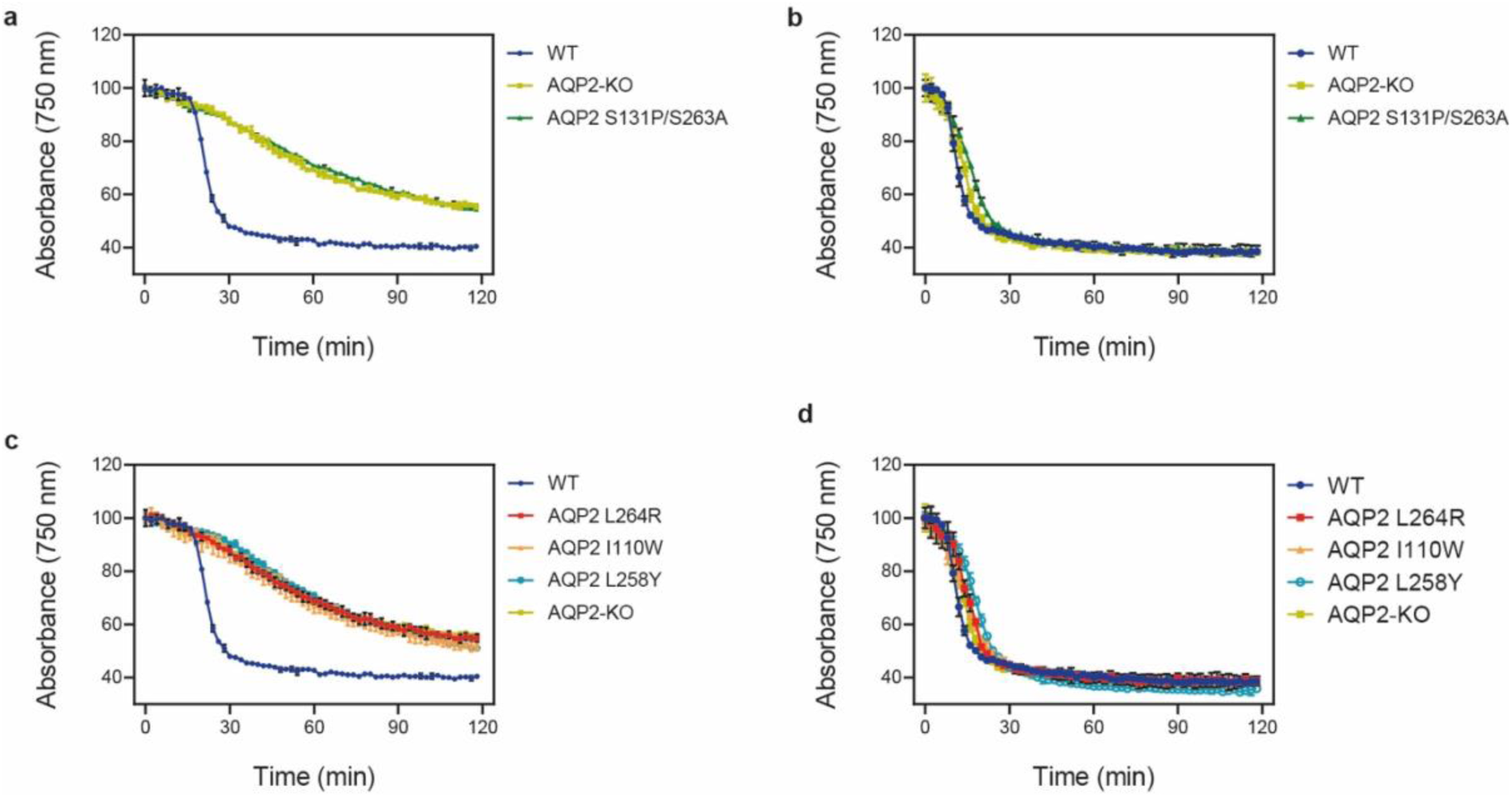
Lysis assay of *T. brucei* cells. **a**-**d,** Lysis assay with *T. b. brucei* wild-type and various AQP2 mutant cell lines treated with arsenical compounds: **a** and **c**, cymelarsan; **b** and **d**, phenylarsine oxide, a different arsenic-containing trypanocide that enters the parasite independently of TbAQP2^8,10,12^ and is thus used as a control for transporter-related arsenic resistance versus resistance to arsenic *per se*. The cells were placed in a cuvette and treated with either compound at t = 10 min. All points shown are the average of triplicate determinations and SD. When error bars are not visible, they fall within the symbol. The slow decline with cymelarsan over time in AQP2-KO and the mutant cell lines is attributable to residual uptake of the compound through the TbAT1/P2 transporter^88,89^.

**Fig. S12.**
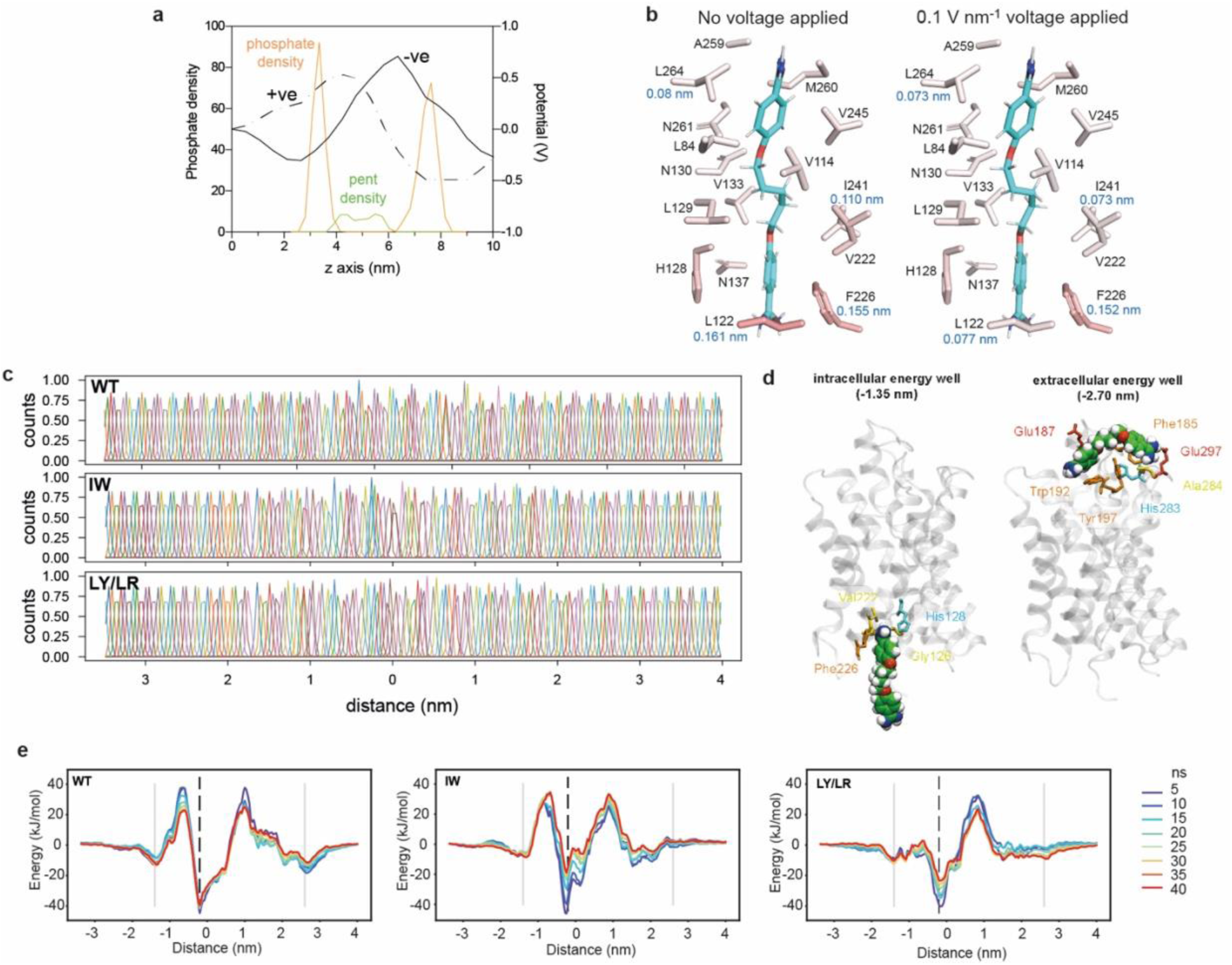
Molecular dynamics simulations. **a**, **Molecular dynamics simulations. a**, Plot of the membrane potential (V) as applied via an electric field, and calculated using the Gromacs tool *gmx potential*. Shown are potentials produced by an electric field applied in the forward (-ve; solid line) or backward (+ve; dashed line) direction. For context, the densities of lipid phosphate atoms and pent atoms are shown as orange and green traces, as measured using *gmx density*. **b**, RMSF calculations were run on monomeric TbAQP2 with either no membrane voltage or a 0.1V nm^-1^ voltage applied (in the physiological direction). Shown are residues in contact with the pentamidine molecule, coloured by RMSF value. RMSF values are shown for residues Leu122, Phe226, Ile241, and Leu264. The data suggest the voltage has little impact on the flexibility or stability of the pore lining residues. **c**, Histograms for the umbrella sampling simulations used to generate the landscapes in Fig. 4d. The data show considerable overlap between windows. **d**, Snapshots taken from post umbrella sampling windows for WT TbAQP2 at -1.35 nm and +2.7 nm. **e**, Conversion plots for the PMF landscapes. Shown are landscapes calculated for increasing lengths of simulations (length in ns). After about 30 ns, the landscapes converged.

**Table S1.**
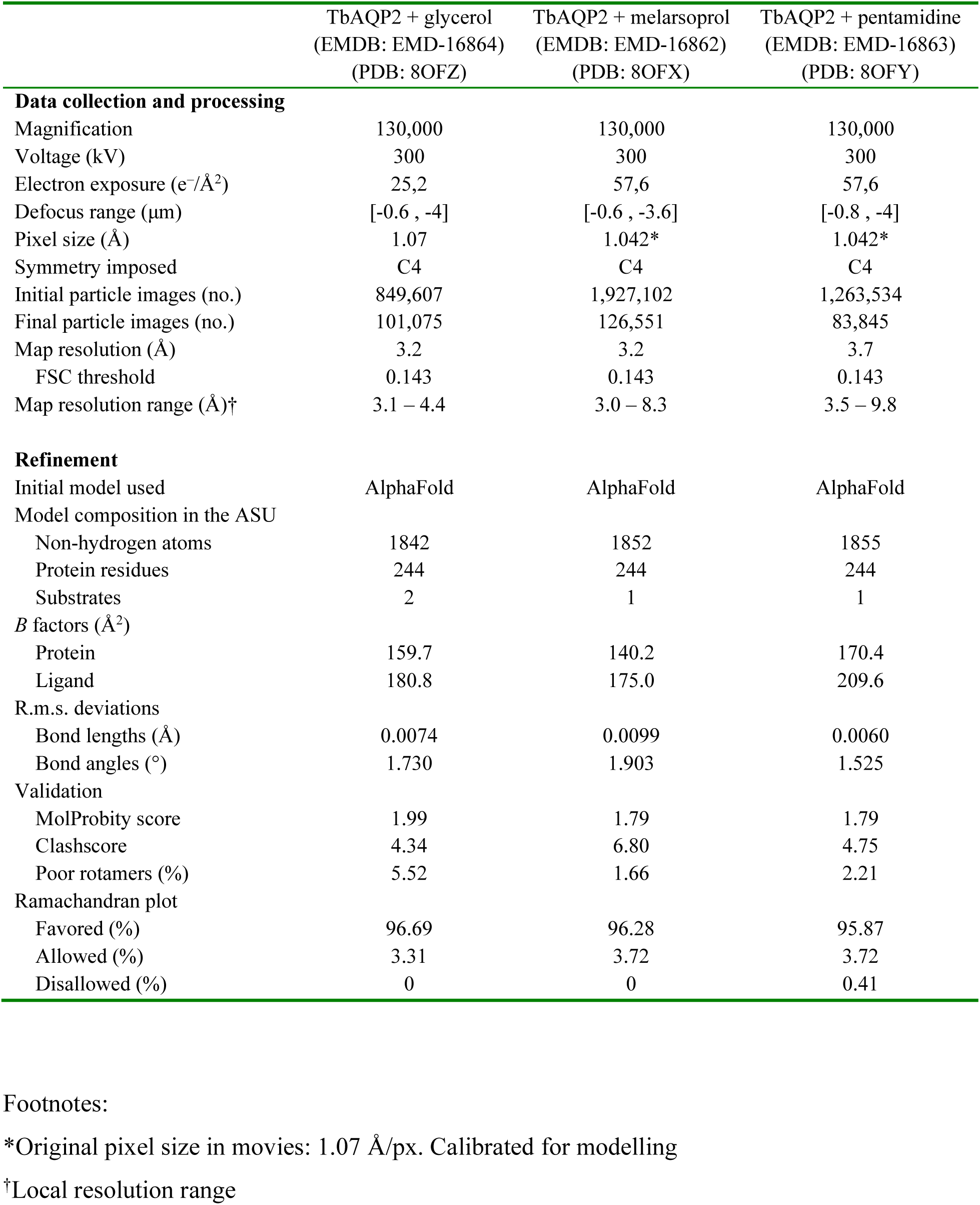
Cryo-EM data collection, refinement and validation statistics.

|  | TbAQP2 + glycerol<br>(EMDB: EMD-16864)<br>(PDB: 8OFZ) | TbAQP2 + melarsoprol<br>(EMDB: EMD-16862)<br>(PDB: 8OFX) | TbAQP2 + pentamidine<br>(EMDB: EMD-16863)<br>(PDB: 8OFY) |
| --- | --- | --- | --- |
| **Data collection and processing** |  |  |  |
| Magnification | 130,000 | 130,000 | 130,000 |
| Voltage (kV) | 300 | 300 | 300 |
| Electron exposure (e<sup>-</sup>/Å<sup>2</sup>) | 25,2 | 57,6 | 57,6 |
| Defocus range (µm) | [-0.6 , -4] | [-0.6 , -3.6] | [-0.8 , -4] |
| Pixel size (Å) | 1.07 | 1.042* | 1.042* |
| Symmetry imposed | C4 | C4 | C4 |
| Initial particle images (no.) | 849,607 | 1,927,102 | 1,263,534 |
| Final particle images (no.) | 101,075 | 126,551 | 83,845 |
| Map resolution (Å) | 3.2 | 3.2 | 3.7 |
| FSC threshold | 0.143 | 0.143 | 0.143 |
| Map resolution range (Å)<sup>†</sup> | 3.1 – 4.4 | 3.0 – 8.3 | 3.5 – 9.8 |
| **Refinement** |  |  |  |
| Initial model used | AlphaFold | AlphaFold | AlphaFold |
| Model composition in the ASU |  |  |  |
| Non-hydrogen atoms | 1842 | 1852 | 1855 |
| Protein residues | 244 | 244 | 244 |
| Substrates | 2 | 1 | 1 |
| *B* factors (Å<sup>2</sup>) |  |  |  |
| Protein | 159.7 | 140.2 | 170.4 |
| Ligand | 180.8 | 175.0 | 209.6 |
| R.m.s. deviations |  |  |  |
| Bond lengths (Å) | 0.0074 | 0.0099 | 0.0060 |
| Bond angles (°) | 1.730 | 1.903 | 1.525 |
| Validation |  |  |  |
| MolProbity score | 1.99 | 1.79 | 1.79 |
| Clashscore | 4.34 | 6.80 | 4.75 |
| Poor rotamers (%) | 5.52 | 1.66 | 2.21 |
| Ramachandran plot |  |  |  |
| Favored (%) | 96.69 | 96.28 | 95.87 |
| Allowed (%) | 3.31 | 3.72 | 3.72 |
| Disallowed (%) | 0 | 0 | 0.41 |

\*Original pixel size in movies: 1.07 Å/px. Calibrated for modelling
<sup>†</sup>Local resolution range

**Table S2.** Details of molecular dynamics simulations on TbAQP2.

| System | Box size | No. atoms | Setup | Simulation length |
| --- | --- | --- | --- | --- |
| AQP2 WT tetramer | 10x10x10 nm | ca. 95,000 | Pentamidine | 5 x ca. 300 ns |
| AQP2 WT tetramer | 10x10x10 nm | ca. 95,000 | Apo | 5 x ca. 300 ns |
| AQP2 WT monomer | 6.5x6.5x10 nm | ca. 45,000 | Pentamidine | 5 x 800 ns |
| AQP2 WT monomer | 6.5x6.5x10 nm | ca. 45,000 | Pentamidine and +ve electric field | 3 x ca. 600 ns |
| AQP2 WT monomer | 6.5x6.5x10 nm | ca. 45,000 | Pentamidine and -ve electric field | 3 x ca. 1400 ns |
| AQP2 WT monomer | 6.5x6.5x10 nm | ca. 45,000 | Pentamidine steered MD | 1 x 5 ns pull |
| AQP2 WT monomer | 6.5x6.5x10 nm | ca. 45,000 | Pentamidine PMF | 167 x 40 ns windows |
| AQP2 WT monomer | 6.5x6.5x10 nm | ca. 45,000 | Apo | 5 x 800 ns |
| AQP2<sub>L258Y_L264R</sub> monomer | 6.5x6.5x10 nm | ca. 45,000 | Pentamidine | 5 x 800 ns |
| AQP2<sub>L258Y_L264R</sub> monomer | 6.5x6.5x10 nm | ca. 45,000 | Pentamidine steered MD | 1 x 5 ns pull |
| AQP2<sub>L258Y_L264R</sub> monomer | 6.5x6.5x10 nm | ca. 45,000 | Pentamidine PMF | 167 x 40 ns windows |
| AQP2<sub>L258Y_L264R</sub> monomer | 6.5x6.5x10 nm | ca. 45,000 | Apo | 5 x 800 ns |
| AQP2<sub>I110W</sub> monomer | 6.5x6.5x10 nm | ca. 45,000 | Pentamidine | 5 x 800 ns |
| AQP2<sub>I110W</sub> monomer | 6.5x6.5x10 nm | ca. 45,000 | Pentamidine steered MD | 1 x 5 ns pull |
| AQP2<sub>I110W</sub> monomer | 6.5x6.5x10 nm | ca. 45,000 | Pentamidine PMF | 167 x 40 ns windows |
| AQP2<sub>I110W</sub> monomer | 6.5x6.5x10 nm | ca. 45,000 | Apo | 5 x 800 ns |

